# The composition of motor recovery revealed by syllable-level behavioral analysis after spinal cord injury

**DOI:** 10.1101/2023.10.31.564826

**Authors:** Jaclyn T. Eisdorfer, Sherry Lin, Joshua K. Thackray, Thomas Theis, Ana Vivinetto, Olisemeka Oputa, Hannah D. Nacht, Alana M. Martinez, Monica Tschang, Malaika Mahmood, Ashley Tucker, Carolin Ruven, Shailee Pusuloori, Lance Zmoyro, Megan V. Phu, Derin Birch, Suneel Kumar, Abraira Lab Computational Group (Collaborative Group/Consortium), Melitta Schachner, Vibhu Sahni, Phillip Popovich, Adam R. Ferguson, Dana McTigue, Vicki M. Tysseling, Jennifer Dulin, Edmund Hollis, Sandeep R. Datta, Victoria E. G. Abraira

## Abstract

The brain-spinal cord axis generates movement by assembling motor primitives into coordinated sequences. Spinal cord injury (SCI) disrupts this neuroaxis, impairing not only locomotion, but the full repertoire of behavior. Traditional scales for quantifying recovery collapse this complexity into predefined locomotor-focused criteria that obscure heterogeneity in recovery. To quantify the full behavioral repertoire following SCI, we adapted motion sequencing (MoSeq) to identify sub-second behavioral “syllables” and capture their usage and sequential organization without predefined features. We identified biomechanically distinct variants within syllable classes that are shared across injury severities and found that recovery is jointly structured by injury severity and individual mouse identity. Changes in sequences, however, unfold along a conserved temporal trajectory. By compressing behavior into a single metric, we uncovered clusters of coevolving locomotor and non-locomotor behaviors. These results frame SCI recovery with repertoire-level changes, where adaptive strategies emerge from constrained access to motor primitives and their sequences.

## MAIN

The brain and spinal cord form a unified axis that generates movement by combining motor primitives into complex behavior^1,2^. Rooted in the principles of ethology, this architecture gives rise to rich behavioral repertoires where actions are flexibly assembled into coordinated sequences^3–8^. Spinal cord injury (SCI) disrupts this neuroaxis by damaging circuitry essential for the expression and coordination of motor primitives^9–11^. As a result, SCI constrains both the set of available actions and the higher-order structure through which those actions can be combined. Emerging work supports this repertoire-wide impact of SCI where recovery-related changes extend across both locomotor and non-locomotor domains^12,13^. Understanding recovery therefore requires quantification of injury-induced influence across the full behavioral repertoire and the temporal sequencing through which that repertoire is organized.

Despite this complexity, recovery after SCI is most commonly evaluated using measures that focus on isolated aspects of movement, particularly those involving locomotion. As a result, large portions of the behavioral repertoire are excluded, including non-locomotor actions (like grooming, rearing, pausing) which are also integral outputs of the motor system and are ethologically central to how animals interact with their environments^14,15^. In preclinical mouse models, this exclusion is reflected in widely used assessments: The field’s standardized ordinal measure of hindlimb motor recovery in mice, the Basso Mouse Scale (BMS)^16,17^, relies on predefined criteria scored by human observers to provide coarse, locomotor-centered descriptions of function. Higher-resolution, automatic pose estimations of isolated movement features used in kinematics analyses can be derived through computer vision tools, but they are likewise limited to features that must be specified in advance^18–20^. Consequently, these locomotor-centric outcome measures do not capture non-locomotor behaviors or how they coevolve with locomotion during recovery, such that animals classified as functionally similar may in fact represent collapsed behavioral strategies.

Beyond individual behaviors, changes in behavioral sequencing can distinguish whether recovery reflects the restoration of preinjury strategies or the emergence of alternative, compensatory solutions under injury-imposed limitations^12,21,22^. Indeed, functional improvement is often accompanied by changes in movement coordination, such as shifts in interlimb timing and context-dependent recruitment of motor patterns^23,24^. However, while temporal-based approaches like gait analyses capture structured recovery dynamics within predefined locomotor contexts, they do not capture how the motor system assembles and transitions between behaviors across the full repertoire. As a result, the broader sequential structure of recovery, such as which actions follow which and how this behavioral syntax organizes during recovery, remains largely unmeasured.

To address these gaps, we examined spontaneous behavior across recovery in mice spanning a biologically meaningful range of SCI severities. We custom adapted a machine-learning framework, motion sequencing (MoSeq)^7,22,25–29^, to segment spontaneous behavior into repeated, sub-second “syllables” (locomotion, rearing, grooming) and capture their usage frequencies and sequential transitions without predefined features. Using this approach, we show that SCI alters behavioral composition across the locomotor and non-locomotor axes and that these changes are shaped by injury severity and mouse individual identity. In contrast, the changes to behavior sequences are constrained primarily by time post-injury, indicating a shared temporal structure to recovery. Syllable-level descriptions uncovered differences among animals assigned identical BMS scores, thus revealing hidden heterogeneity as an emergent and measurable feature of recovery. They also provided improved predictive access to underlying injury neurobiology as compared to conventional BMS scores. Compression of repertoire-wide behavior into a single metric uncovered clusters of coevolving locomotor and non-locomotor behaviors. These findings position recovery following SCI as repertoire-level changes under a disrupted brain-spinal cord axis, where adaptive strategies emerge from constrained access to motor primitives and their flexible assembly into coordinated sequences.

## RESULTS

### Spinal cord injury reorganizes the spontaneous behavioral repertoire beyond locomotor recovery

To understand how SCI reshapes behavior across recovery, we examined spontaneous mouse behavior across a biologically meaningful gradient of injury severities that sample overlapping spinal and supraspinal circuits and a broad range of functional capacities^30,31^. Mice received lower thoracic Transection (Tx), Moderate (Mod, 50 kDyn) or Mild (30 kDyn) contusions, Sham laminectomy, or no surgery (Naive) and were recorded before surgery (preinjury), 2 days post-operation (DPO), and weekly for 70 days (**Fig. 1a**). We first examined raw movement patterns to determine whether broad behavioral patterns differ longitudinally across injury severities. Because of sex-dependent differences^32–35^, male mice are shown in the main figures and female mice in Extended Data. At 2 DPO, raw movement paths were visibly stratified by severity (**Fig. 1b**) and group-level quantification across all groups and timepoints confirmed that injury reduced the percentage of time spent moving (**Fig. 1c, Extended Data Fig. 1a**). These findings indicate that injury imposes constraints on spontaneous behavior.

**Fig 1.**
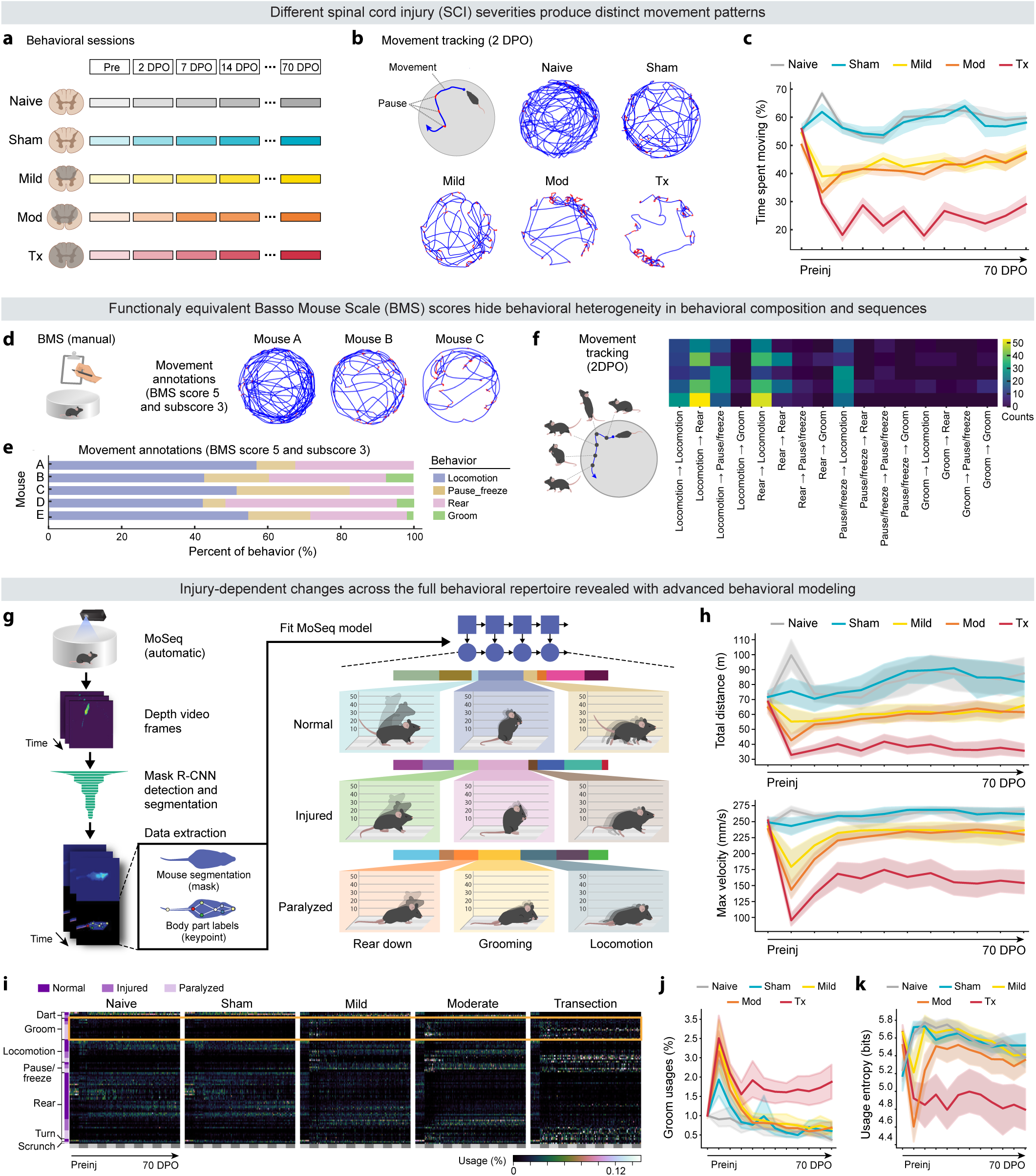
Spinal cord injury reorganizes spontaneous behavior beyond locomotor recovery. a, Experimental setup. Individual mice (n = 71 male) were recorded in an open-field arena to assess their mobility pre- and post-operation. Sessions included: preinjury (“pre”), 2 days post operation (DPO), 7 DPO, and weekly thereafter. Groups included: (no-SCI) Naive and Sham; and (SCI) 30 kDyn Mild and 50 kDyn moderate (Mod) contusions, and transection (Tx). **b,** Representative raw movement trajectories at 2 DPO reveal stratification of injury severity. **c,** Quantification of spontaneous movement by group over time. Data were analyzed by ANOVA and Dunnett’s multiple comparison for c, h, and j-k (*P* values can be found in Source Data 1). **d,** Raw movement trajectories from mice assigned the same functional equivalence as described by the BMS illustrate different spontaneous movements. **e,** Manual ethograms of BMS-matched animals reveal individualized behavioral compositions. **f,** Transition matrices derived from manually annotated ethograms of BMS-matched animals show individualized behavioral sequencing patterns. **g,** We adapted an established unsupervised decomposition framework (MoSeq^7,29^) for analysis of the full behavioral repertoire. Complex behavior was segmented into sub-second behavioral “syllables" spanning different hindlimb engagements. **h,** Total distance traveled (top) and maximum velocity (bottom) by group over time. **i,** Behavioral summary heatmap of syllables demonstrates distinct behavior compositions across groups during recovery (93 syllables identified, Supplemental Table 1). Example grooming behaviors (orange box) highlights group and timepoint differences. **j,** Quantification of grooming behavior differences shows that a non-locomotor behavior changes significantly across different injury severities during recovery. **k,** Behavioral composition entropy extrapolated from all syllable usages. Higher entropy indicates a more diverse range of available behaviors. BMS, Basso Mouse Scale; MoSeq, motion sequencing. Data and code used for analyses underlying all figures are available at: https://doi.org/10.5281/zenodo.18014529.

We next examined whether injury also introduced individualized constraints on behavior that were not resolved at the group level by analyzing behavior across animals that achieved the same functional milestone under the field’s gold standard metric, BMS. BMS is an ordinal, observer-based scoring system that evaluates hindlimb locomotor recovery based on criteria like weight support, coordination, and trunk stability^16^. To examine behavior within a BMS-derived functional milestone, we focused on our most frequently observed BMS score-subscore combination: 5-3. Despite identical BMS assignment, we found that animals displayed qualitatively different raw movement patterns (**Fig. 1d**). To uncover what might be driving these differences, we manually annotated behavior, which revealed individualized frequencies of locomotion, pausing/freezing, rearing, and grooming behaviors (**Fig. 1e**). Transition matrices of these manually annotated ethograms showed animal-specific switching structures (**Fig. 1f**). Thus, equivalent BMS scores can summarize non-equivalent behaviors, which may suggest animals rely on individualized behavioral solutions including and outside of BMS criteria.

To resolve recovery-related differences in behavioral strategies, we required a high-resolution, unbiased approach capable of capturing the full spontaneous repertoire to longitudinally compare animals with different injury severities. We adapted an unsupervised behavioral decomposition framework (motion sequencing, MoSeq) that represents behavior as sequences of repeated, sub-second behavioral “syllables” (rear, locomotion, groom, etc.) and quantifies their usage frequencies and temporal organization^7,22,25–29^. Importantly, standard MoSeq segmentation is optimized for animals with normal hindlimb weight-bearing and therefore cannot capture low-lying trunk and hindlimb movements common after SCI. To overcome this limitation, we modified the pipeline to incorporate a Detectron2-based multi-task Region-Based Convolutional Neural Network (R-CNN)^36^ for full body segmentation post-SCI (**Fig. 1g, Supplemental Fig. 1**, see Methods). Validation of adapted MoSeq readouts using traditional open-field metrics recapitulated the severity-associated stratification observed in raw movement tracking (**Fig. 1h**, **Extended Data Fig. 1b**), confirming that MoSeq preserves gross behavioral signatures.

We then ask whether differences in all behaviors, including those not typically assessed during recovery but captured by MoSeq, reflected genuine biological phenomena (**Fig. 1i**, **Extended Data Fig. 1c**). We trained separate male and female MoSeq models and identified over 90 unique syllables in each model (**Supplemental Tables 1-2**). We found that syllables outside the classically scored locomotor axis showed changes that aligned with prior work. For example, we observe that grooming-like behaviors (referred to hereafter as “grooming” syllables) are enriched in paralyzed animals, gradually diminish over time during recovery, and remain elevated in animals with permanent paralysis (**Fig. 1j**, **Extended Data Fig. 1d**). This observation is consistent with prior studies on excitotoxic SCI and the onset of mechanical allodynia, which have shown that animals experiencing hypersensitivity following SCI display elevated grooming^37,38^. To capture these repertoire-wide shifts quantitatively, we computed a diversity index (entropy), which revealed that injury decreases behavior availability (**Fig. 1k**, **Extended Data Fig. 1e**). These results suggest that non-locomotor behaviors also participate in the injury-induced behavioral repertoire, as opposed to emerging as incidental byproducts.

### Injury severity and individual identity jointly structure the behavioral recovery landscape

Consistent with the fact that different SCI severities damage overlapping neural circuitry^30,31^, we hypothesized that distinct injuries may also share subsets of behaviors. To characterize this, we first dimensionally reduced syllable usage features from all behavioral sessions across severities and timepoints. This embedding organized along a continuum from paralysis to normality, with graded motor deficits in between and notable overlap among groups (**Fig. 2a**, **Extended Data Fig. 2a**). Within this space, more severe injuries produced larger and more enduring displacements from preinjury behavioral composition, whereas mild and sham injuries returned more rapidly toward baseline (**Fig. 2b**, **Extended Data Fig. 2b**, **Supplemental Fig. 2ab**). This pattern suggested that different combinations of injury severity and time post-injury converge onto overlapping regions of behavioral space. Linear classifiers confirmed this convergence by distinguishing groups better at early timepoints when motor deficits were most pronounced and declining in performance as behavioral representations overlapped across groups (**Fig. 2c**, **Extended Data Fig. 2c**). These observations highlight that the source of this shared structure across injury severities may reside at the level of the syllables themselves.

**Fig 2.**
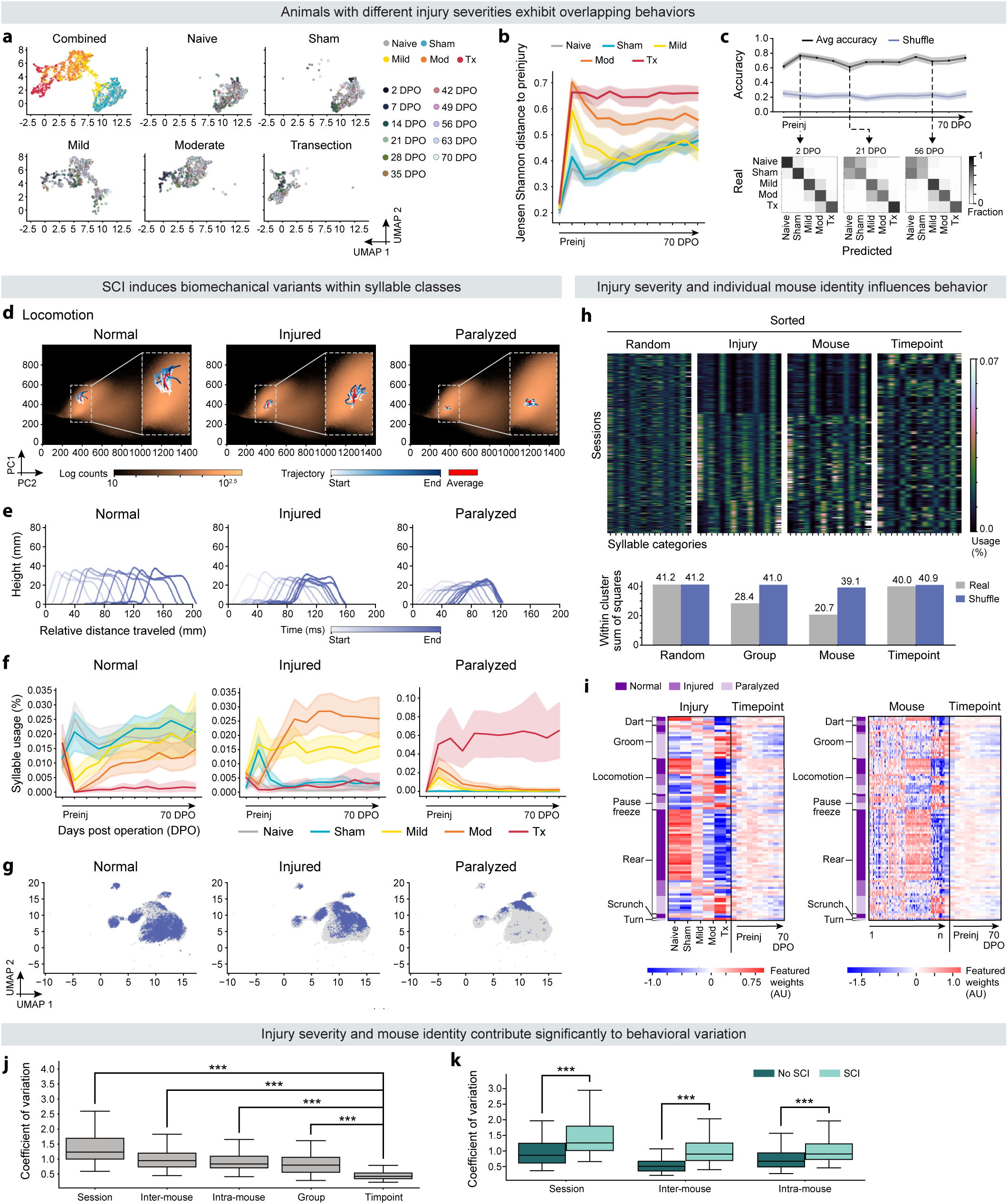
Injury severity and individual identity jointly structure exhibited behaviors during recovery. a, UMAPs of syllable usages. Data points from different timepoints overlap within and between groups, indicating shared behaviors across injury severities. **b,** Jensen Shannon distance to preinjury state (ANOVA with Dunnett’s multiple comparisons test for b and f; n=71; *P* values for Fig. 2 can be found in Source Data 3). **c,** Linear classifier accuracy in distinguishing groupwise syllable usages across timepoints (top). Example confusion matrices (bottom), where darker shading indicates better performance and lighter shading indicates confusion. **d,** Projection of example locomotion behaviors into PC embedding space shows conserved within-syllable trajectories distinct embedding regions correspond to normal, injured, and paralyzed biomechanical variants. **e,** Spinograms, a visualization of animal height and relative distance across syllable duration, further illustrate biomechanical differences among variants. **f,** Longitudinal usage of example locomotion syllable variants. **g,** UMAP embeddings of weighted behavioral sequences, colored by locomotor syllable variants. **h,** Behavioral composition heatmaps sorted randomly or by group, mouse, or timepoint (top). Within cluster sum of squares (bottom) describes purity within sorting, with a smaller number denoting more structure. **i,** Weights from generalized linear models with syllable usage as a function of: injury types and timepoints (left) and mouse ID and timepoints (right). **j,** CV of syllable usages across: all sessions (session), different mice (intermouse), sessions within a mouse (intramouse), groups (group), and timepoints (timepoints) (for j and k: unpaired t-test; *n*=71; \*\*\**P* < 0.001; *P* values can be found in Source Data 3). **k,** Same as j, but split by no SCI vs. SCI reveals that SCI contributes more to observed variation. Mod, moderate; Tx, transection; DPO, days post operation; PC, principal component; UMAP, Uniform Manifold Approximation and Projection; CV, coefficient of variation.

The presence of shared syllables raised the possibility that recovery does not rely on entirely novel behaviors, but instead involves conserved actions expressed through altered motor strategies. To test this, we hierarchically clustered syllables based on similarities of their autoregressive dynamics (see Methods). Video inspection confirmed that many post-SCI syllables resembled preinjury syllables, but were executed with altered posture and reduced hindlimb engagement. Together, this indicated persistence of preinjury behavioral motifs in biomechanically modified forms. Principal Component Analysis (PCA)^39^ embeddings revealed that within each behavioral class (locomotion, groom, rear, etc.), syllables separated into three biomechanical variants distinguished primarily along PC1, which encoded animal height (**Fig. 2d**, **Extended Data Fig. 2d**, **Supplemental Fig. 2cd**). The body posture of these variants differed in normal, intermediate, and low height off the ground (**Fig. 2e**, **Extended Data Fig. 2e**). Functionally, recovery was accompanied by decreased usage of intermediate- and low-height variants and increased usage of normal-height variants (**Fig. 2f**, **Extended Data Fig. 2f**). Despite their shared motif identity, syllable class variants participate in different higher-order sequencing structures (**Fig. 2g**, **Extended Data Fig. 2g**, **Supplemental Fig. 2ef**). To capture these functional distinctions in a form amenable to quantitative comparison across injury severities and recovery, we grouped syllable variants into three functional categories (**Supplemental Tables 1-2**): "normal syllables" are observed preinjury, by the Naive and sham groups, and later in recovery in a subset of the mild and moderately injured animals; “injured syllables” denote weightbearing movements that emerge during recovery and are absent in non-injured animals, such as unstable locomotion, and likely reflect compensatory strategies adopted in response to injury-induced physical constraints; and "paralyzed syllables" refer to behaviors that occur when an animal drags its hindlimbs.

These conserved functional variants prompted us to ask what governs their expression across recovery. It is well established that recovery after SCI is strongly constrained by injury severity40; however, increasing evidence indicates that animals also exhibit inter-individual variability that reflects meaningful differences in how animals adapt to injury^26,41,42^. To assess the relative contributions of these factors, we sorted syllable usage heatmaps by injury severity, mouse individuality, and timepoint. Sorting by injury severity and mouse identity produced distinct, organized patterns, whereas sorting by timepoint failed to yield a clear structure (**Fig. 2h**, **Extended Data Fig. 2h**). Generalized linear models supported that behavioral composition was more strongly influenced by injury severity and individual mouse identity (**Fig. 2i**, **Extended Data Fig. 2i**). Quantification of explained variation further showed that injury severity and mouse identity contributed significantly more to behavioral differences than timepoint, with this effect being most pronounced in injured animals (**Fig. 2jk**, **Extended Data Fig. 2jk**). These findings suggest that while injury severity is one of dominant factors that influences post-SCI behavior^40^, individual variability introduces meaningful differences that reflect how each mouse uniquely adapts to injury.

### Behavioral sequencing reorganization follows a time-dependent structure across recovery

Behavior is not solely defined by individual syllables, but also by the way these syllables are organized and expressed in sequences to produce fluid behavior^7,22,25–28^. Following SCI, transitions in behavior have been extensively studied in the context of gait, with prior research demonstrating how injury-induced alterations disrupt gait patterns and transitions between gaits, as well as how these changes evolve throughout recovery^43–48^. However, whether SCI reorganizes the broader structure of behavioral sequencing beyond locomotion, and whether this reorganization depends on injury severity or other factors, remains unclear. As such, we next characterized the probabilistic structure of behavior as it changes throughout recovery^22^, where “transition probabilities” reflect the likelihood of the animal switching from one syllable to the next^25^. We quantified averaged absolute changes in transition probabilities to capture how sequences reorganize (**Fig. 3a**). Similar to syllable usages (**Fig. 2**, **Extended Data Fig. 2**), the entropy of transition probabilities scaled with motor function, where animals with greater motor function had higher transition entropy (**Fig. 3b**, **Extended Data Fig. 3a**). This suggests that higher functional capacity is associated with more flexible and diverse behavioral sequencing, whereas greater motor deficits constrain the range of permissible behavioral transitions and lead to more stereotyped and predictable syllable sequences.

**Fig 3.**
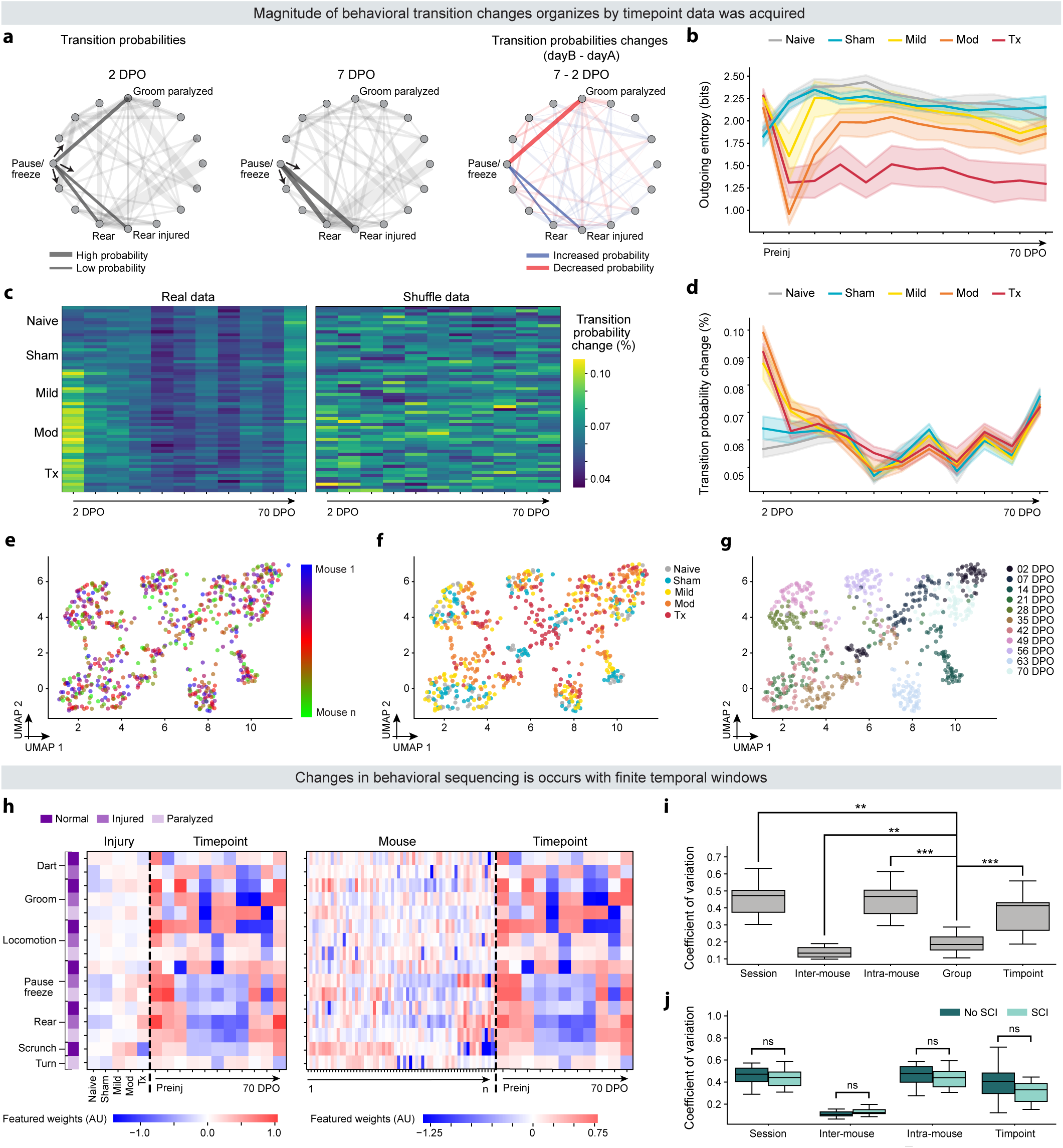
Behavioral sequence reorganization follows a time-dependent structure across recovery. A, Illustrative example of syllable transitions evolving over time, with visualizations provided for 2 DPO (left), 7 DPO (middle), and the difference in transitions between 7 DPO and 2 DPO (right). **b,** Average syllable outgoing entropy over time for different groups over time. Higher entropy reflects a broader range of possible next behaviors, whereas lower entropy indicates more predictable and constrained sequencing (ANOVA with Dunnett’s multiple comparisons test for b and d; n = 71; *P* values can be found in Source Data 5). **c,** Heatmaps of average absolute changes in outgoing transition probabilities over time using real (left) and shuffle (right) data. **d,** Quantification of average absolute outgoing transition differences in c shows that the magnitude of sequence reorganization decreases with recovery. **e-g,** UMAP embeddings of transition change vectors across syllable categories, colored by mouse (e), injury severity (f), or timepoint (g). Transition changes cluster predominantly by timepoint, whereas weaker organization is shown if grouped by mouse of injury. **h,** Weights from generalized linear models relating changes in transition probabilities to injury severity and timepoint (left) or mouse identity and timepoint (right). Stronger weights associated with timepoint indicate a temporal influence on the organization of behavioral sequences. **i,** CV of syllable usages was analyzed across various groupings: session (within all sessions), inter-mouse (across different mice), intra-mouse (within a single mouse), group (across different groups), and timepoints (across different timepoints) (for j and k: unpaired t-test; *n*=71; \*\**P* < 0.01; \*\*\**P* < 0.001; *P* values can be found in Source Data 5). **j,** Same as i, but split by no SCI vs. SCI did not reveal significant differences between the two groups. Mod, moderate; Tx, transection; UMAP, Uniform Manifold Approximation and Projection; CV, coefficient of variation.

To characterize how this sequencing reorganizes across recovery, we examined the magnitude of changes in transition probabilities. Interestingly, transition probability changes were mostly similar across groups within the same timepoint (**Fig. 3c**, **Extended Data Fig. 3b**). A notable exception was observed immediately post-operative, where the largest overall changes in motor behavior occurred across injured groups. This acute phase reflects the expected disruption caused by the injury itself^16,30,31^, including changes in exhibited syllables used during this period. Across all groups, the magnitude of transition changes diminished steadily with time, indicating the stabilization and habituation of behavior as time progressed^49–52^ (**Fig. 3d**, **Extended Data Fig. 3c**). These results suggest that despite profound differences in motor impairment, there is a shared, time-dependent structure to behavior sequence organization across all groups.

Such temporal conservation could imply that recovery is constrained by when behavioral sequences can reorganize, rather than allowing unrestricted sequence flexibility. To assess the structure of these constraints, we analyzed transition-change vectors using dimensionality reduction analyses, which revealed that transition changes predominantly clustered based on timepoint (**Fig. 3e-g**, **Extended Data Fig. 3d-f**). Generalized linear models corroborated this result, showing that changes in transition probabilities were more strongly influenced by timepoints post-SCI than by other factors (**Fig. 3h**, **Extended Data Fig. 3g**). A comparison to usage changes further supported results (**Supplemental Fig. 3**). Quantification of variation analyses showed a dominant temporal influence (**Fig. 3i**, **Extended Data Fig. 3h**), with no observed significant differences between SCI and no-SCI groups (**Fig. 3j**, **Extended Data Fig. 3i**). This finding underscores that observed changes were primarily influenced by time-dependent factors rather than injury status. As such, temporal rules may bound behavioral sequence flexibility and raise the possibility that there may be “permission windows” during which behavioral transitions are most malleable.

### Repertoire-level behavioral analysis reveals hidden heterogeneity across animals and bidirectional changes in behavior during recovery

Behavioral scales for SCI provide standardized summaries of functional status that enable cross-laboratory comparisons and therapeutic evaluation. In mouse SCI models, BMS is the field’s gold standard due to its reproducibility, sensitivity, and ability to predict lesion severity^16^. By design, BMS is restricted to predefined, human-visible locomotor criteria; however, our findings demonstrate that recovery also involves coordinated changes across non-locomotor behaviors. To evaluate whether repertoire-level descriptions provide a more complete account of functional status, we first assessed their concurrent and predictive validities53 with BMS criteria. To test concurrent validity, we examined if syllables could encode BMS criteria. Pearson’s correlations demonstrated strong, directionally appropriate relationships to BMS criteria, including strongly positive correlations to normal syllables and strongly negative correlations to paralyzed syllables (**Fig. 4ab**, **Extended Data Fig. 4ab**, **Supplemental Fig. 4a**). Injured syllables showed positive correlations with partial recovery criteria (like plantar stepping), but weaker, negligible, or even negative correlations with higher-level recovery criteria (like coordination). Furthermore, despite reliance on a single downward-facing camera view that does not explicitly visualize some criteria scored by BMS, such as paw placement, we found that syllables could accurately predict all BMS criteria (**Fig. 4cd**, **Extended Data Fig. 4cd**). This indicates that syllable-based descriptions capture human-observed determinants of locomotion16,17,54 as unique biomechanical signatures.

To further test concurrent validity, we next asked whether repertoire-level descriptions resolve additional, behaviorally meaningful differences within BMS-defined functional status. We examined this by summarizing syllable usages using principal component analysis (PCA), which situates each animal-day according to its overall behavioral composition. When PCA embedding plots were color-coded by BMS scores and subscores, substantial dispersion was apparent among animals assigned the same functional score (**Fig. 4e**, **Extended Data Fig. 4e**). Comparing the behaviors associated with specific BMS score-subscore to the syllables representing the same scores, we observed markedly greater behavioral variability in the syllable space (**Fig. 4fg**, **Extended Data Fig. 4fg**). Quantifying this variation confirmed that many behavioral solutions exist at each BMS score-subscore combination (**Fig. 4hi**, **Extended Data Fig. 4hi**). These results expose hidden heterogeneity among animals assigned identical BMS scores and potential inconsistencies between adjacent scores^55^, which may reflect biologically meaningful differences in compensatory strategies or circuit level adaptations that are not resolvable through the locomotor behavioral axis alone.

**Fig 4.**
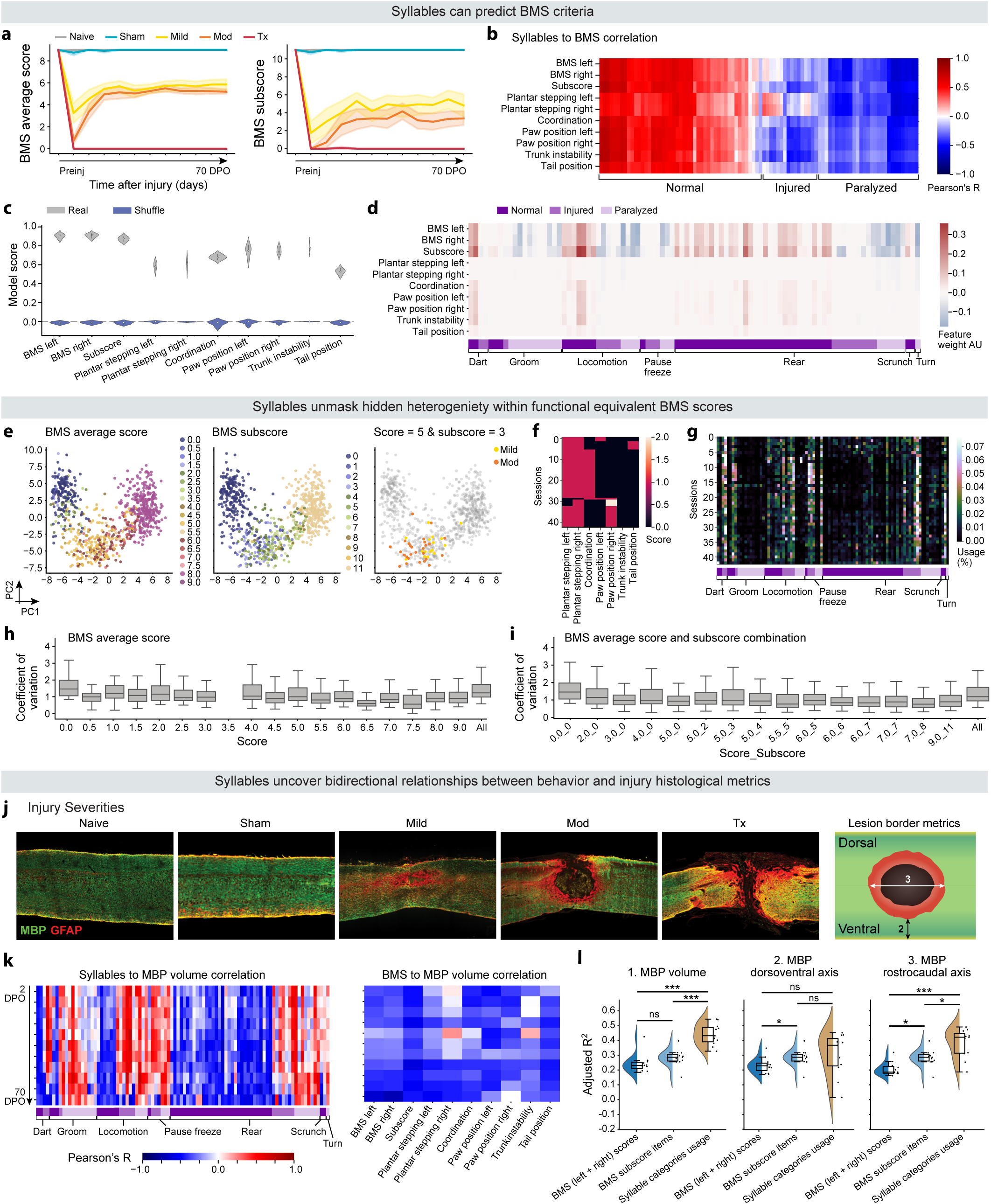
Quantitative behavioral descriptions reveal hidden heterogeneity among animals with equivalent locomotor function. a, BMS average scores (left) and subscores (right) (ANOVA with Dunnett’s multiple comparisons test; n = 71; *P* values can be found in Source Data 7). **b,** Pearson’s correlation between syllable usages and BMS demonstrate directionally appropriate relationships. **c,** Cross validations of regularized linear regression models predicting BMS criteria from syllable usages. **d,** Feature weights from BMS prediction models in c, where darker colors indicate syllables that are important for predicting each BMS score. **e,** PCA computed from syllable usages colored by BMS scores (left), BMS subscores (middle), and animals with a BMS score-subscore combination of 5-3 (right) reveals dispersion among animals assigned equivalent functional scores. **f,** Heatmap of BMS criteria for animals with a score-subscore combination of 5-3. **g,** Corresponding heatmap of syllable usages for the same animals as f demonstrates the higher resolution afforded by repertoire-based descriptions in distinguishing behavioral strategies masked by BMS. **h-i,** CV of syllable usages within individual and across all BMS scores (h) and across all BMS score-subscore combinations (i) reveals variability within nominally equivalent functional states. **j,** Representative sagittal spinal cord sections at the lesion epicenter across injury severities and schematic illustration of lesion metrics. **k,** Pearson’s correlations for injury-induced demyelination volumes and syllable usages (left) and BMS criteria (right). Syllables exhibit bidirectional relationships with lesion metrics, whereas BMS criteria show uniformly negative correlations. **i,** Adjusted R^2^ values from correlation models of lesion metrics with BMS Scores, BMS subscores, and syllable category usages for all timepoints (unpaired t-test; n = 71; \**P* < 0.05, \*\**P* < 0.01, \*\*\**P* < 0.001; *P* values can be found in Source Data 7). Mod, moderate; Tx, transection; BMS, Basso Mouse Scale; PCA, principal component analysis; MBP, myelin basic protein; GFAP, glial fibrillary acidic protein.

Given the strong influence of lesion burden on motor outcomes^54^, we next tested the predictive validity^53^ of syllable-based descriptions relative to BMS by examining associations between syllable usage and total lesion volume and the spatial dimensions of the injury along the dorsoventral and rostrocaudal axes at the lesion epicenter^56^ (**Fig. 4j**). Pearson’s correlations revealed dynamic, bidirectional relationships between syllable usages and lesion metrics across recovery, with individual syllables exhibiting positive or negative correlations with lesion metrics that varied in strength across recovery (**Fig. 4k left**, **Extended Data Fig. 4j left**, **Supplemental Fig. 3b-e top**). This indicates that injury severity is reflected not only in the gain of behaviors associated with improved motor performance, but also the suppression of behaviors associated with greater motor deficits. In contrast, BMS criteria exhibited uniformly negative correlations with lesion metrics with minimal temporal variation (**Fig. 4k right**, **Extended Data Fig. 4j right**, **Supplemental Fig. 3b-e bottom**), reflecting a predominantly monotonic measure of functional improvement^17^. To quantify the predictive implications of this bidirectionality, we used regression models to compare syllable-based and BMS-based relationships to lesion characteristics. Syllables achieved significantly better model fits across all timepoints for total injury volume and lesion metrics along the rostrocaudal axis, and comparable performance to BMS criteria along the dorsoventral axis (**Fig. 4k**, **Extended Data Fig. 4k**). These findings suggest that bidirectional behavioral representations across the full repertoire provide improved access to lesion characteristics. Such coordinated organization across sets of syllables also indicates that groups of behaviors may coevolve during recovery.

### A unified metric of the aggregated behavioral repertoire reveals coevolving clusters of syllable usages across the paralysis-to-normality continuum

If recovery is organized by coordinated, bidirectional changes across subsets of behaviors, then determining whether such coevolving structure exists, and which behaviors coevolve together, requires a common functional axis through which behavioral trajectories can be compared across animals. To this end, we asked whether the full behavioral repertoire could be aggregated into a continuous metric analogous to BMS, while retaining repertoire-level resolution. We applied PCA to syllable usage features to yield a low-dimensional embedding that organized behavior along a continuum from paralysis to normality, with graded levels of motor deficits in between (**Fig. 5a**, **Extended Data Fig. 5a**). To quantify an animal’s position along this continuum, we applied diffusion-based inference using the Palantir approach^57,58^, which produced a scalar metric we termed the “recovery score” that summarized each animal’s functional status based on its repertoire-wide behavioral composition (**Fig. 5bc**, **Extended Data Fig. 5bc**, see Methods). Recovery score curves (**Fig. 5d**, **Extended Data Fig. 5d**) show strong concordance with classical BMS curves (**Fig. 4a**, **Extended Data Fig. 4a**), indicating that recovery scores are consistent with BMS in principle and share a similar behavioral semantic representation. Importantly, recovery scores retained reversible interpretability in terms of constituent behaviors, including positive correlations with distance traveled and maximum velocity and negative correlations with grooming usage (**Fig. 5e**, **Extended Data Fig. 5e**). To evaluate whether this preserved behavioral meaning translated into improved functional quantitation and relationships with injury-related structure, we employed regression models to compare the first PC of the behavioral embedding with BMS and recovery scores. Recovery scores exhibited smaller and more evenly distributed residuals for recovery scores, indicating a better linear model fit (**Fig. 5fg**, **Extended Data Fig. 5fg**). Further, recovery scores outperformed BMS scores and subscores in correlating with total injury volume and rostrocaudal axis distribution, while showing comparable performance along the dorsoventral axis (**Fig. 5h**, **Extended Data Fig. 5h**). This is consistent with syllable-level analyses (**Fig. 4l**, **Extended Data Fig. 4k**), indicating that aggregation of the behavioral repertoire preserves the necessary granularity to retain behavior–lesion relationships.

**Fig 5.**
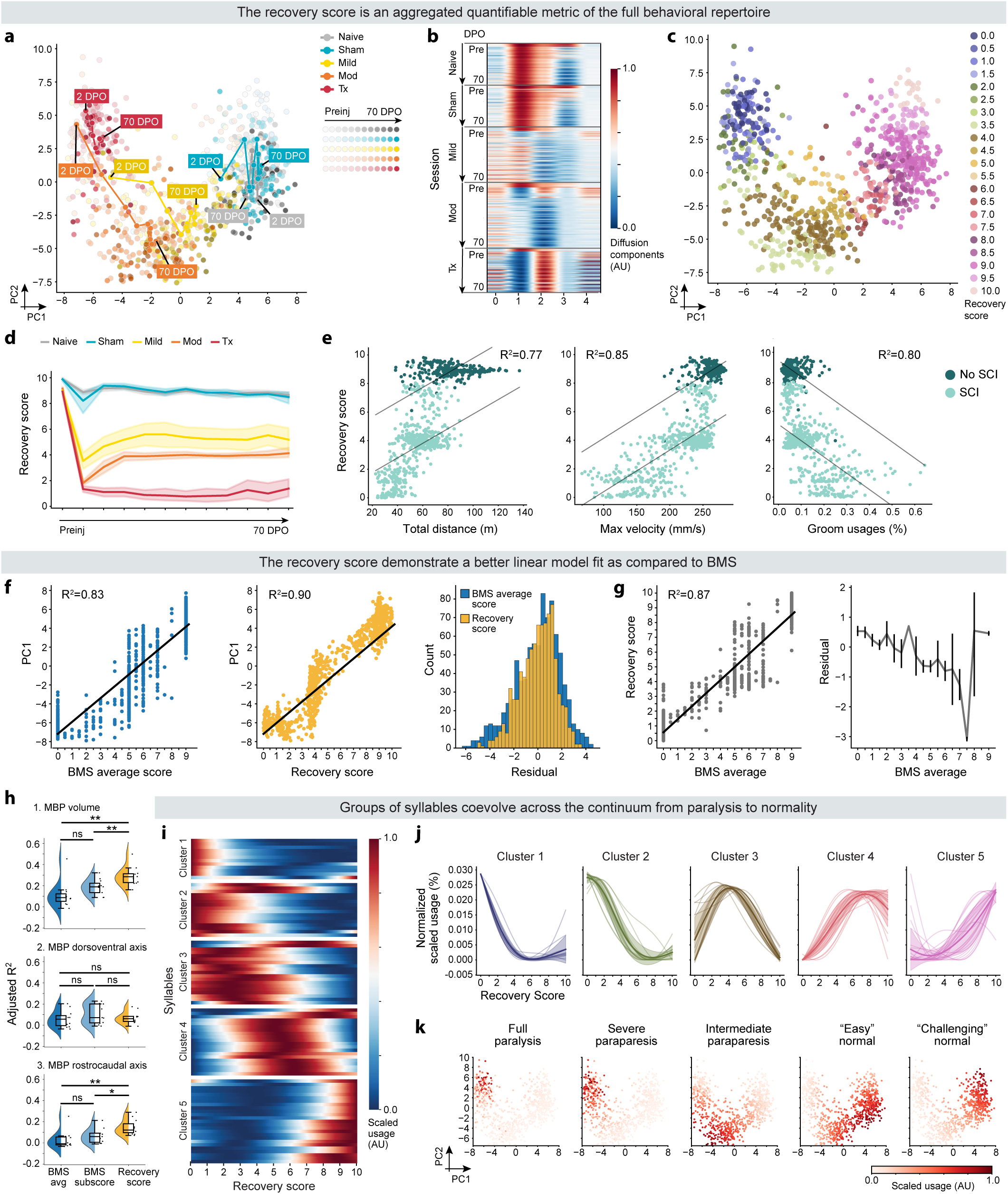
A repertoire-derived recovery score reveals clusters of coevolving behaviors across the paralysis-to-normality continuum. a, PCA embedding of syllable usage colored by group, shaded by timepoint, and labeled with each group’s averaged longitudinal behavior. **b,** Diffusion components derived from syllable usages summarize variation in behavior across recovery and provide the basis for computing the “recovery score.” **c,** Projection of recovery scores onto the PCA embedding. **d,** Recovery Scores recapitulate expected severity-dependent recovery curves (ANOVA with Dunnett’s multiple comparisons test; n = 71; *P* values can be found in Source Data 9). **e,** Correlations between recovery scores and total distance traveled, maximum velocity, and groom usages reveal strong linear relationships (Ordinary Least Squares linear regression for e-g). **f,** Comparison of linear model fits between behavioral PC1 with BMS scores (left) and recovery scores (middle). Residual values (right) demonstrate that recovery scores are more uniformly distributed. **g,** Correlations between BMS scores and recovery scores (left). Model residuals indicate variability in how BMS aligns with recovery scores (right). **h,** Adjusted R^2^ values from correlation models relating lesion metrics with BMS scores, BMS subscores, and recovery scores (unpaired t-test; n=71; \**P* < 0.05; *P* values can be found in Source Data 9). **i,** Heatmap of syllable usages across recovery scores. Leiden clustering discovered groups of syllables that coevolve within the recovery score space. **j,** Normalized syllable usages within each syllable cluster. Thin lines denote individual syllables, bold lines indicate cluster means, and shading represents standard deviation. **k,** PCA embeddings colored by normalized mean cluster-specific syllable usages, where darker shades represent higher usage. Clusters were semantically labeled based on functionally interpretable behavioral regimes (Supplemental Table 3). Mod, moderate; Tx, transection; BMS, Basso Mouse Scale; PCA, principal component analysis; MBP, myelin basic protein.

With a shared functional axis established, we asked whether the bidirectional changes in behavior observed across recovery arise from independent modulation of individual behaviors or from coordinated, coevolving subsets of behaviors. To distinguish between these possibilities, we applied Leiden clustering^59^ to syllable usage trends, which identified groups of syllables with similar usage patterns (**Fig. 5ij**, **Extended Data Fig. 5ij**, **Supplemental Fig. 5c-f**, see methods). Mapping cluster-averaged syllable usage onto the PCA embedding showed alignment with recovery trajectories over time and along the recovery score axis (**Fig. 5k**, **Extended Data Fig. 5k**). Group-wise analyses of syllable usages, velocities, and heights across clusters validated these functional distinctions (**Supplemental Fig. 5gh**). These results suggest that recovery after SCI may be organized by coordinated, multi-behavioral clusters that correspond to different functional states.

To understand the functional demands that may distinguish these clusters, we characterized and semantically labeled each cluster according to its behavioral composition (**Supplemental Tables 3-4**): Full Paralysis, Severe Paraparesis, Intermediate Paraparesis, "Easy" Normal, and "Challenging" Normal. The first three clusters encapsulated canonical post-SCI behaviors^16^: Full Paralysis was marked by hindlimb dragging; Severe Paraparesis involved major deficits with some hindlimb movement; and Intermediate Paraparesis was characterized by weight-bearing deficits and noticeable instability. Interestingly, we also observed two clusters with all “normal” behaviors: the Easy Normal cluster included behaviors that might be less physically demanding^60,61^, such as wall-assisted rears and slow locomotion, and were accessible to animals with mild SCI at later recovery stages. In contrast, the Challenging Normal cluster, observed exclusively in animals without SCI, was comprised of behaviors that require greater hindlimb engagement and coordination^62^, such as unassisted high rears and fast locomotion or darting. The emergence of these distinct clusters may indicate that recovery after SCI reflects selective access to functionally achievable subsets of behavioral solutions under physical constraints related to injury severity.

## DISCUSSION

The brain-spinal cord axis flexibly assembles motor primitives across a rich behavioral repertoire into coordinated sequences that enable animals to adaptively interact with their environment^1–4^. SCI disrupts the spinal and supraspinal pathways along this axis, thereby constraining which actions are available and how those actions can be temporally organized^12,13^. Despite this repertoire-wide disruption, recovery following SCI has traditionally been evaluated using locomotor-centric assessments, such as BMS^16^, which necessarily compress behavior into human-derived coarse, ordinal scores. By quantifying behavior across the full repertoire, we show that SCI alters both locomotor and non-locomotor behaviors and reorganizes the sequential structure through which available actions are assembled. More specifically, recovery was characterized by a bidirectional architecture in which increases in adaptive behaviors were accompanied by suppression of deficit-associated behaviors. This reciprocal pattern aligns with prior observations across locomotor kinematics^23^, skilled reaching^63^, and grooming-related phenotypes^37^, where functional improvements coincide with attenuation of maladaptive movements. Framing recovery as a bidirectional process thus positions functional restoration milestones as a simultaneous acquisition and suppression of behaviors.

Across the full behavioral repertoire, injury severity emerged as a dominant driver of behavioral composition^40^, but individual differences also exerted a non-negligible influence. Intrinsic factors, including an animal’s baseline preinjury behaviors and individual adaptive capacity, may play a role in shaping recovery^26,41,42^. Such individuality may arise from circuit-level properties, including differences in baseline excitability^64^, sensorimotor integration^65,66^, and action-selection biases^28,67^. This individuality likely also contributes to our observation that animals assigned the same BMS score and subscore do not express the same underlying behaviors. This hidden heterogeneity suggests that similar functional performance can arise from distinct behavioral solutions, a phenomenon previously documented in kinematics analyses^68,69^. These findings support two possible interpretations: identical scores may correspond to similar recovery states expressed through distinct behavioral strategies or may obscure substantive differences in functional restoration.

Beyond behavioral composition, the organization of behavioral sequences provides additional insight into recovery. In intact systems, brain-spinal cord interactions coordination transitions between motor actions^70,71^. However, circuit disruptions, such as those that occur due to SCI, can alter transition structure and may result in potential maladaptive reorganization^12,22^. Surprisingly, we observed a consistency in the degree of change within sequences across groups at a given timepoint, regardless of injury severity or even the presence of an injury. This temporal consistency suggests the existence of a finite, measurable ceiling effect that limits how much sequential reorganization can occur within a given interval. Such a ceiling may reflect biomechanical constraints of the neuromuscular system, finite rates of circuit adaptation, or time-dependent stabilization of behavior^60,72,73^. In injured systems, this ceiling may highlight a finite capacity for behavioral sequence reconfiguration within defined temporal windows^60^ and raises intriguing questions about whether conserved organizational principles govern sequence adaptation in both injured and uninjured systems.

To compare recovery across mice, we compressed the behavioral repertoire into a unified numeric metric (the “recovery score”) that is functionally analogous to BMS^16^, but retained both locomotor and non-locomotor behaviors. These multi-domain behaviors organized into distinct functional clusters, a finding consistent with prior work on the restitution of multiple motor functions after SCI, such as improvements in bladder and sexual function accompanying gains in leg movement^74,75^. Importantly, the existence of clusters that contain both locomotor and non-locomotor behaviors underscore the need for a more precise scoring system, as manually scored traditional recovery assessments alone cannot capture the complexity of SCI recovery. While this study offers a promising foundation for developing such an approach, several limitations remain, including constraints imposed by a downward-facing camera view, technical inability to accommodate varying mouse sizes, and the purposeful focus on bilateral contusion SCI models whose behavior spanned paralysis to normalityss^30,31^. Incorporating broader injury models, mouse strains, treatment conditions, and complementary kinematics approaches (like Keypoint-MoSeq^76^) will be required to further refine the recovery score and evaluate its potential as a broadly applicable recovery assessment. Future research could also explore how the emergence of specific behaviors within clusters aligns with neural activity patterns and how these patterns differ between clusters.

In conclusion, our study posits that recovery after SCI is defined by coordinated, coevolving changes across both locomotor and non-locomotor behavioral domains. This coevolution indicates that non-locomotor behaviors are integral to recovery and may reflect shared neurobiological mechanisms with locomotor behaviors. By quantifying the full spontaneous behavioral repertoire, this work revealed a previously largely unmeasured dimension of recovery, with the goal of shifting toward recovery analyses that jointly consider locomotor and non-locomotor behaviors.

## MATERIALS AND METHODS

### Subjects

Male and female 8- to 9-week-old C57BL/6 (Strain Code 027) mice (n = 71 male, n = 68 female) were procured from Charles River Laboratories (Wilmington, MA, USA). The mice were provided ad libitum access to food and water and were maintained in a 12-hour light and 12-hour dark cycle within the animal facility at the Division of Life Sciences, Nelson Biology Laboratories, Rutgers University. Spinal cord injury (SCI) surgeries were conducted on these mice at an age of 9-10 weeks. All housing, surgical procedures, behavioral experiments, and euthanasia were carried out in strict adherence to Rutgers University’s Institutional Animal Care and Use Committee (IACUC) guidelines, with the experiments following the ARRIVE guidelines.

### Surgical Procedures

The mouse thoracic SCI contusion model was conducted following the protocol described previously^16,77^. Briefly, 9- to 10-week-old mice were anesthetized with isoflurane (NDC 66794-017-25, Piramal Critical Care, Inc., Bethlehem, PA, USA), administered at 5% initially and reduced to 2% during surgery. A thermo-regulated heating pad was utilized to monitor and sustain the body temperature throughout the surgery. Prior to surgical incision, the upper-back skin was shaved and sanitized using alternating preparations of Betadine scrub (NDC 67618-151-32, Purdue Products L.P., Stamford, CT, USA) and 70% ethanol. Preemptive and post-operative analgesia were provided by injecting 0.1 ml of 0.125% Bupivacaine (NDC 0409-1163-18, Hospira, Lake Forest, IL, USA) subcutaneously around the incision site. The slow-release Buprenorphine formulation, Ethiqa XR (Fidelis Pharmaceuticals, LLC, North Brunswick, NJ, USA), was subcutaneously administered at 3.25 mg/kg.

To prevent corneal drying, lubricant ophthalmic ointment (Artificial Tears, NDC 59399-162-35, Akorn Animal Health, Lake Forest, IL, USA) was applied to the eyes post-anesthesia induction. A 3 cm skin incision was made along the median line on the animal’s back, followed by laminectomy at the vertebrae level halfway between T9 and T10. The contusion SCI was induced using the IH Infinite Horizon Impactor (Precision Systems and Instrumentation, LLC; model IH-0400). The mouse was secured on the experimental table using spinal cord stabilizing forceps, and the exposed spinal cord was precisely positioned. The impactor tip was placed over the center of the exposed spinal cord. A force of 30 kDyn was used for mild injuries, and a force of 50 kDyn was employed for moderate injuries, both with a dwell time of 0 ms. The muscles were then sutured in two separate anatomical layers (Suture 4-0 Polyglactin 910 FS-2, Cat. No. 6549052, Henry Schein, Melville, NY, USA) and the incision closed with wound clips (Reflex-9, 9 mm, Cat. No. 201-1000, Cell Point Scientific, Inc.,Gaithersburg, MD, USA). Sham-injured animals underwent laminectomy but did not receive a contusion.

Animals classified as "naive" received drugs during the approximate duration of the surgery but were not incised. Animals designated for complete transections underwent transections following laminectomy using aspiration and sectioning with microscissors as previously described^16,30^ using great care to ensure complete severance of all axon fibers.

### Postoperative monitoring and care

Following surgery, mice were individually housed in cages with white Alpha Dri bedding (W.F. Fisher & Son, Branchburg Township, NJ, USA) to monitor for potential bladder infections and bleeding. Subsequently, mice received subcutaneous injections of cefazolin (25 mg/kg) and 0.5 ml saline once daily for three days after injury. Additionally, cages were positioned partially on 37°C thermo-regulated heating pads for the duration of the study. Monitoring of food intake was performed using Labdiet Rodent Diet 5001 pellets (Cat. No. LD5001, W.F. Fisher & Son) placed within the cages, with DietGel (Clear H2O Dietgel 76A, Cat. No. CH2ODG762OZ, Fisher & Son) provided if necessary. Daily assessments for dehydration were conducted, and HydroGel (Clear H2O Hydrogel, Cat. No. CH2OHG2OZ) was administered as needed along with supplemental saline injections. Bladder care was administered twice daily from the day after injury until euthanization to prevent urine retention. Cases of hematuria were treated with Baytril (2.7 mg/kg) over a 10-day period. Mice were withdrawn from the study if they experienced weight loss exceeding 80% of their initial body weight or exhibited severe or persistent autophagia.

### Behavior: Basso Mouse Scale (BMS)

At 2 days and 1-10 weeks post-spinal cord injury, locomotor recovery in mice was assessed using the BMS^16^. Mice were placed in the center of a small wading pool (open field) with a 44-inch diameter for 4 minutes, and hind limb movements were evaluated by two trained scorers on a scale ranging from 0 to 9. A score of 0 indicated no hind limb movement, while higher scores denoted improved hind-limb joint coordination, with 9 representing "normal" walking. In cases of discrepancy between scorers, the lower score was selected. If necessary, the sub-score table, considering parameters such as plantar stepping, coordination, paw position, trunk instability, and tail position, was utilized.

### Behavior: Raw movement tracking and manual ethograms

On the day of acquisition, mice were transferred to the behavior room for a 30-minute habituation. RGB data was acquired using a Microsoft Kinect v2 camera, pointed downwards towards a circular arena, connected to a computer running the program kinect2-nidaq (v0.2.4alpha). Individual mice were gently placed in the arena, and their naturalistic behavior over a 20-minute period was recorded. Captured video frames were each 512 x 424 px, collected at 30 Hz. Open-field videos were analyzed using Noldus EthoVision XT17^78^. For raw movement tracking, automatic animal detection settings were used with minor parameter adjustments as needed to ensure continuous tracking of each mouse across the entire arena, including during rearing along the arena walls, for the 20 min recordings. A smooth tracking curve was applied to stabilize the centerpoint of the mouse during tracking. Manual ethograms were generated for the first five minutes of recordings from five mice assigned a BMS score–subscore of 5–3. Each annotated behavior (locomotion, rear, groom, pause/freeze) was assigned shortcut keys and annotated by a trained observer. Subsequent analyses of manually annotated behavioral composition and transition structure were performed using the same software.

### Behavior: MoSeq Framework

Here and below, we employed methodology as previously described^7,22,25,26,26–28^ and customized it as required.

### Data acquisition

On the day of acquisition, mice were transferred to the behavior room for a 30-minute habituation. Raw depth data was acquired using a Microsoft Kinect v2 camera, pointed downwards towards a circular arena, connected to a computer running the program kinect2-nidaq (v0.2.4alpha). Individual mice were gently placed in the arena, and their naturalistic behavior over a 20-minute period was recorded. Captured depth video frames were each 512 x 424 px, collected at 30 Hz, with each pixel encoded as a 16-bit unsigned integer indicating the distance of that pixel from the Kinect camera sensor in millimeters. Data was saved in binary format and transferred to another computer system for later processing.

### Data preprocessing

Because injured mice often face paralysis, resulting in overall lower-to-the-ground body positions, we found it difficult to reliably segment mice from the noise and background using the classical computer vision approaches deployed in the conventional moseq2-extract pipeline. Therefore we developed a custom deep-learning-based extraction program, which outputs results that are compatible with the existing MoSeq ecosystem of tools. We adapted the Detectron2 framework released by Facebook AI Research^36^, based on the state of the art mask-R-CNN architecture, as it could jointly propose segmentation masks and keypoint estimations. We use a ResNet 50 FPN backbone which feeds a Region Proposal Network (RPN), and both then feed a series of modular ROI Heads, namely Instance Bounding Box, Instance Segmentation Mask, and Instance Keypoint Regression. We generated a training set of 4,529 background-subtracted depth images, split into training (4,076 images) and test (453 images) subsets, which we annotated with masks and eight keypoints: nose, left/right ear, neck, left/right hip, tail base, and tail tip. During model training, we applied randomized augmentations including random rotation, random scaling, random brightness, random contrast, random gaussian white noise, and random particle noise (to simulate dust particles in the acquisition environment). We trained the model for 100,000 iterations, at which point performance plateaued with the following COCO performance metrics: bbox (AP: 64.812, AP50: 97.202, AP75: 73.423), keypoints (AP: 45.851, AP50: 93.768, AP75: 37.795), and segm (AP: 55.680, AP50: 98.673, AP75: 56.266).

For utilizing this model in extraction of real-world videos, we constructed a parallelized processing pipeline that would 1) read raw depth video frames from saved files, remove background and non-ROI pixels, 2) submit images for deep learning inference producing estimates of boxes, masks and keypoints, 3) post process these features and generate cropped and rotated mouse images and masks (in the same style as classical extraction), 4) generate a preview video visualization of the extraction results and save results in HDF5 format in a way that mimics the outputs produced by classical extraction. This whole process is able to extract a twenty minute depth video in about twelve minutes on a computer system equipped with a AMD Ryzen 7 3800X (8-Cores @ 3.89 GHz), 64 GB RAM, Nvidia RTX 2080 SUPER, and network attached storage (1Gbps link).

### Behavioral Modeling and Analysis

The preprocessed data underwent dimensionality reduction and modeling through the MoSeq pipeline^25^, resulting in frame-by-frame descriptions of the animal’s dynamics (i.e., velocity, position) and syllable labels. Syllable usage fractions were computed across all sessions for analysis. Semantic labels were manually assigned to syllables by examining their associated dynamics and behaviors in the recorded movies. Noise syllables were excluded from further analysis, resulting in 92 syllables for male mice (“male model”) and 93 syllables for female mice (“female model”). Behavioral diversity within a session was quantified using usage entropy, defined as 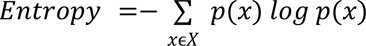, where 𝑥 represents the usage fraction of each syllable within the session, and 𝑋 is the usages of all syllables. Behavioral similarity between sessions was assessed using the Jensen–Shannon distance, calculated with the scipy.spatial.distance.jensenshannon function.

Transitions between syllables are characterized by outgoing transition probabilities, which represent the likelihood of one syllable transitioning into another. The diversity of these syllable transitions were quantified using outgoing entropy (same formula as above), where *x* represents the probabilities of one syllable transitioning into each other syllable in the session. To analyze changes in outgoing probabilities (Fig 3), syllables were aggregated into 17 syllable categories, so that outgoing probabilities are aggregated to represent one group of syllables transitioning to another group of syllables (i.e., from dart to pause, from locomotion to rear, etc.). The mean absolute differences in outgoing probabilities between two consecutive timepoints within each mouse were computed to capture transitions. As a control, outgoing syllable categories were shuffled within each mouse to generate a “shuffle control”.

Classical locomotion metrics, such as total distance traveled and maximum velocity, were derived from MoSeq outputs. Distance traveled was calculated as the sum of the Euclidean distances between mouse centroid coordinates across two consecutive frames. Maximum velocity was determined as the 99 percentile of session velocities as a means of excluding extreme values from noise.

### Behavioral Distance Analysis

MoSeq behavioral distance is a metric used to assess the similarity between two syllables. Behavioral distance was calculated as previously described^28^ using the function “get_behavioral_distance()” from the moseq2-viz library using default parameters. Briefly, the function computes the correlation between the 3D pose trajectories associated with each syllable, which are encapsulated by the autoregressive process that models the two syllables. The behavioral distance is then derived by subtracting the correlation value from one. To analyze patterns in the behavioral distance matrix, hierarchical clustering was applied using a Euclidean distance metric and average linkage method. This clustering allowed for the grouping of syllables based on their behavioral similarity, which then facilitated downstream analysis of behavioral structure and patterns.

### Predicting BMS Scores from Syllable Usages

Correlations between BMS features and syllable usages were computed using ‘scipy.stats.pearsonr()‘ for each pair of syllables and BMS metrics. Correlation R-values were plotted using seaborn.

Regression models to predict BMS metrics given syllable usages were constructed using scikit-learn. Values were first standardized using ‘StandardScalar()‘ and then fed to an ‘ElasticNet‘ model with the parameter ‘l1_ratiò set to 0.1. A 5-fold cross-validation strategy that balanced injury types and timepoints across CV splits was employed for model training and evaluation. Shuffle models were also generated, in which the syllable usage values were randomly shuffled prior to being fed to the model. Model performance, reported as the coefficient of determination (R^2^) and measured on the held-out portion of data, across the distribution of CV splits was plotted as violins using seaborn. Model coefficients, averaged across the CV splits, were plotted as a heatmap using seaborn.

### Syllable PC Trajectory

Syllable trajectories in the PC space were computed and visualized from the PC scores generated during the dimensionality reduction step in the MoSeq analysis pipeline. To smooth the trajectories, the PC scores were smoothed using a Gaussian kernel with a window length 5. The background of the visualization featured a binned 2D histogram to represent the density of points in PC space. To illustrate syllable dynamics, the syllable segments with the most frequent occurrence of same-length trajectories were selected to increase the count of representative examples. Trajectories originating from similar starting points in the PC space and exhibiting the closest shape in terms of trajectory dynamics were chosen to show the syllable dynamics in the PC space.

### Syllable sequence Uniform Manifold Approximation and Projection (UMAP)

Syllables were labeled with semantic labels such as “locomotion,” “rear,” etc., and subsequently converted into one-hot encoding where a value of 1 indicated the animal is performing the syllable and 0 indicates it is not.

Each row of the resulting dataframe corresponds to a single frame of behavior. To incorporate contextual information about temporally adjacent syllables, a Gaussian kernel was applied to the one-hot encoded dataframe, thereby allowing each row to reflect syllables performed in close temporal proximity. Each point on the UMAP represents a single frame with the UMAPs thus capturing structural patterns in syllable sequences over time.

### UMAPs of syllable usages and degree of syllable category transition changes

UMAP was employed to reduce the dimensionality of the data for visualizing both syllable usages across groups and timepoints, as well as the degree of syllable category transition changes. For syllable usages, the syllable usage data from each session was z-scored before applying UMAP. The UMAP parameters for this analysis were set as: n_components=2, n_neighbors=20, metric=’euclidean’, and min_dist=0.1. For syllable category transition changes, the mean absolute changes across all syllable categories between two consecutive timepoints for each mouse were z-scored prior to UMAP analysis. The UMAP parameters for this analysis were set as: n_components=2, n_neighbors=20, metric=’cosine’, min_dist=0.3.

### Linear classifiers

Linear classifiers were employed to predict group labels based on syllable usages across all sessions within each timepoint (**Fig. 2g and Ext. Data Fig. 2g**). Model accuracy was calculated from the results of 10 independent restarts of model runs. To ensure dataset balance, the input data was randomly resampled to equalize the number of sessions per group and standardized prior to modeling. A linear support vector classifier (SVC) was utilized, with a Stratified K-Fold (n_split=5) cross validation approach applied to maintain proportional representation of each group across folds. Accuracy was calculated as the ratio of true predictions to total predictions (True Labels/All Labels) and compared against results from shuffled data within each timepoint.

### Within Cluster Sum of Squares (WCSS)

The WCSS was used to quantify the variability within each group, where lower values represent more similarity within the group, and higher values represent more dispersion within the group. To calculate WCSS for each type of sorting, the centroid of each group was computed by averaging the syllable usages across all sessions within the group for each syllable category. Then the sums of Euclidean distances between all sessions and their respective group centroids were computed. The sum of these distances within each group provided group-specific WCSS, which was then subsequently summed across all groups to produce the final WCSS metric. The calculated WCSS values were compared against those derived from data with shuffled group labels to evaluate their significance of the observed clustering.

### General linear model (GLM)

GLMs were applied to analyze the relationship between factors such as group, mouse identity, and timepoint with syllable usage and transition probability changes. For syllable usage, GLMs were constructed using one-hot encoded factors for each session as predictors to model log-transformed, z-scored syllable usage. The models were: syllable usages ∼ group + timepoint; usages ∼ mouse + timepoint. For absolute outgoing transition probability changes, GLMs were similarly designed to predict log-transformed, z-scored average absolute outgoing transition probability changes based on one-hot encoded factors for each mouse at each timepoint. The models were: mean absolute change ∼ group + timepoint; mean absolute change ∼ Mouse + Timepoint. A linear regression model (ElasticNet with α=0.01) was fit to model log-transformed, z-scored, session based, syllable usage for each.

### Coefficient of Variation (CV)

The CV was calculated as the ratio between the standard deviation to the mean and is used to assess the variation of data within specific groupings. In Fig 2J, session CV for each syllable was computed across all sessions; for Inter-mouse, CV for each syllable is computed across mean syllable usages of each mouse; for intra-mouse, CV for each syllable is computed across mean syllable usages of each mouse in each timepoint; for injury, CV for each syllable is computed across mean syllable usages of each injury in each timepoint; for timepoint, CV for each syllable is computed across mean syllable usages of each timepoint in each mouse.

The “No SCI vs. SCI” analysis followed a similar methodology, but the data was divided into two groups: no SCI (Naive, Sham) and SCI (Mild, Mod, Tx). In Fig 3I, the CV for each syllable category was computed across mean absolute outgoing probability changes instead of mean syllable usages and Fig 3J was split by no SCI and SCI. In Fig 4IJ, CV for each syllable was computed across mean syllable usages in each BMS average score and BMS average score+subscore combination.

### Lesion metrics regressions

Linear regressions (ordinary least square) were utilized to predict lesion metrics (MBP volume, MBP along the dorsoventral axis, and MBP along the rostrocaudal axis) at each timepoint. For Fig 4m, the models were: lesion metric ∼ BMS left + BMS right; lesion metric ∼ BMS subscore items; lesion metric ∼ MoSeq syllable categories. Similarly, for Fig 5h, the models were: lesion metric ∼BMS avg score; lesion metric ∼ bms subscore; lesion metric ∼ recovery score. The R^2^ for each model was compared across timepoints using t-tests.

### Recovery scores and syllable usage clusters across different recovery scores

Recovery Scores and syllable usage clusters were derived using a cell differentiation algorithm, Palantir (REF), adapted to capture the progression of recovery from paralysis to normal function. Recovery Scores were calculated from pseudotime (as described in the Palantir algorithm) to represent what stage a session is at in the spectrum from full paralysis to normality. Briefly, the algorithm first applied PCA to reduce the dimensionality of syllable usages across all sessions. The dimensionality-reduced data are used to compute the diffusion maps^58^ to capture developmental trends with parameters set to n_components=5 and knn=30.

Sessions were ordered along the diffusion components by calculating the shortest path between user-defined preinjury normal sessions (fully functional) and 2 DPO sessions (least functional) to assign pseudotime for each session that represent the relative distance to the predefined preinjury normal session. The user-defined sessions that were selected were sessions with the closest Euclidean distance to the respective group syllable means (i.e., the preinjury session with the closest distance to preinjury means, the 2 DPO session in Naive with the closest distance to 2 DPO Naive means, etc.). The Recovery Score was computed as 𝑅𝑒𝑐𝑜𝑣𝑒𝑟𝑦 𝑆𝑐𝑜𝑟𝑒 = (1 − 𝑃𝑎𝑙𝑎𝑛𝑡𝑖𝑟 𝑃𝑠𝑒𝑢𝑑𝑜𝑡𝑖𝑚𝑒) × 10 to scale values to a range between 0 and 10, analogous to theBMS scale. Lower Recovery Scores correspond to reduced motor function, while higher scores reflect more normal motor functions. For visualization, Recovery Scores were rounded to the nearest 0.5 (**Fig. 5c and Extended Data Fig. 5c**). Syllable usage trends along Recovery Scores were analyzed using Mellon Function Estimator fit to 500 evenly spaced numbers between 0 and 10, similar to the palantir.presults.compute_gene_trends function^79^. Nearest neighbor (n_neighbors=35) and Leidesn algorithm are used to cluster syllables with similar trends across the recovery scores. To visualize the cluster’s mean usage hotspots on the PCA plots (**Fig. 5k and Extended Data Fig. 5k**), mean syllable usages are scaled with a MinMaxScalar so that the highest usage is 1 and lowest usage is 0.

### Immunohistochemistry

Following the final behavioral tests at 10 weeks post-SCI, mice were deeply anesthetized with 5% isoflurane (NDC 66794-017-25, Piramal Critical Care, Inc., Bethlehem, PA, USA), followed by transcardial perfusion with saline containing 10 U/mL heparin (Cat No. H3393-100KU, Sigma-Aldrich, St. Louis, MO, USA), and 4% paraformaldehyde (Cat No. 15714-S, Electron Microscopy Sciences, Hatfield, PA, USA). The spinal cord columns were dissected, and the ventral side of the spinal cord was exposed. The tissue was postfixed in 4% paraformaldehyde overnight at 4°C, subsequently washed three times with phosphate-buffered saline (PBS), and stored in PBS containing 0.01% sodium azide for further processing.

For immunohistochemistry of the lesion site, 1 cm long spinal cords containing the lesion site at the center were cryoprotected by overnight immersion in 4°C PBS with increasing sucrose concentrations (10%, 15%, 20%), beginning with the lowest sucrose concentration overnight in each solution. Finally, spinal cords were incubated overnight with 50% Optimal Freezing Temperature (OCT) (Tissue-Tek, Cat No. 25608-930, VWR, Radnor, PA, USA) medium and 50% 20%-sucrose dissolved in PBS. Following these saturation steps, the tissue was frozen in OCT medium, and 20 µm thick serial sagittal cryosections were prepared, mounted on Diamond® White Glass Charged Microscope Slides (Globe Scientific Inc, Cat. No. 1358W), and stored at -20°C for immunostaining. The slides were thawed and dried for 1 hour at 20°C, washed three times with PBS, and then blocked with 3% bovine serum albumin (BSA) (Cat No. RLBSA50, IPO Supply Center, Piscataway, NJ, USA), 10% Normal Goat Serum (NGS) (Cat No. 01-6201, Fisher Scientific, Waltham, MA, USA), and 0.3% Triton-X 100 (X100-100ml, Sigma-Aldrich, St. Louis, MO, USA) for 1 hour at 20°C. Subsequently, the tissue sections were co-stained overnight at 4°C with a primary chicken antibody against the astrocyte marker Glial Fibrillary Acidic Protein (GFAP) (1:1000 in blocking solution; Cat No. 200-901-D60, VWR, Radnor, PA, USA) and a primary rat antibody against myelin basic protein (MBP) (1:50 in blocking solution; Cat. No. MAB386, Sigma-Aldrich, St. Louis, MO, USA). Following five 5-minute washes at 20°C with PBS, the tissue was incubated for 2 hours at 20°C with corresponding Alexa Fluor 488 (1:500, anti-Chicken coupled to Alexa Fluor 488, Cat No. A11039, Fisher Scientific, Waltham, MA, USA) and Alexa Fluor 565 (1:500, anti-rat coupled to Alexa Fluor 546, Cat No. A11081, Fisher Scientific, Waltham, MA, USA) secondary antibodies diluted in blocking solution. The tissue was washed five times for 5 minutes at 20°C with PBS, incubated for 10 minutes with DAPI solution diluted 1:5000 in PBS, and the sections were finally mounted under glass coverslips with Fluoromount-G (Cat No. 100241-874, VWR, Radnor, PA, USA).

### Image acquisition and analysis

Quantitative analysis of the sagittal sections was performed using the automated microscope IN Cell Analyzer 6000 (GE Healthcare, Chicago, IL, USA). Four microscope slides were loaded into the system, and the image capturing protocol from the manufacturer was implemented. The tissue sections were precisely located via preview, and high-resolution images were acquired for subsequent analysis.

## Supporting information

Statistics for all Figures

