## Extended Data Figures for "The composition of motor recovery revealed by syllable-level behavioral analysis after spinal cord injury"

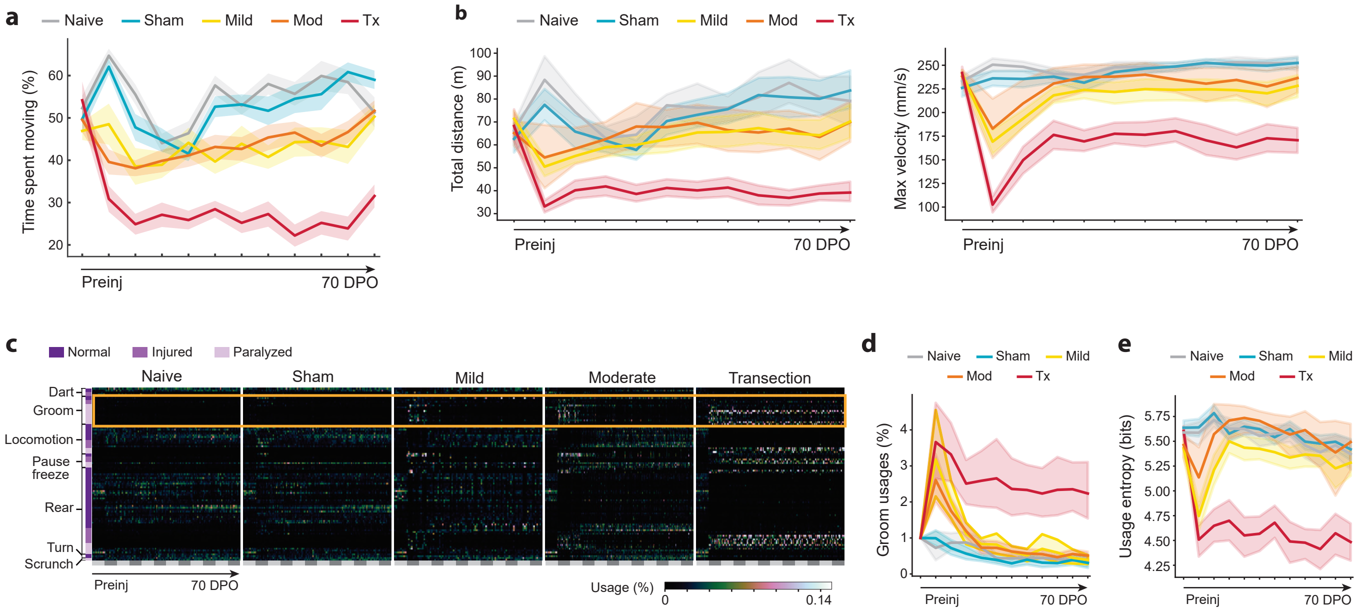


**Extended Data Fig 1. Spinal cord injury reorganizes spontaneous behavior beyond locomotor recovery (female model). a,** Quantification of spontaneous movement by group over time (n=68; ANOVA with Dunnett's multiple comparison test for a, b, d, and e; *P* values can be found in Source Data 2). Same as Fig. 1c, but for females. **b,** Total distance traveled (left) and maximum velocity (right) by group over time. Same as Fig. 1h but for females. **c,** Behavioral summary heatmap with syllables ordered by type and injury severity (95 syllables identified, Supplemental Table 2). Same as Fig. 1i but for females. **d,** Grooming behaviors (aggregated from all grooming syllables usages) by group over time. Same as Fig. 1j but for females). **e,** Behavioral composition entropy by group over time extrapolated from all syllable usages. Same as Fig 1k but for females. Mod, moderate; Tx, transection.

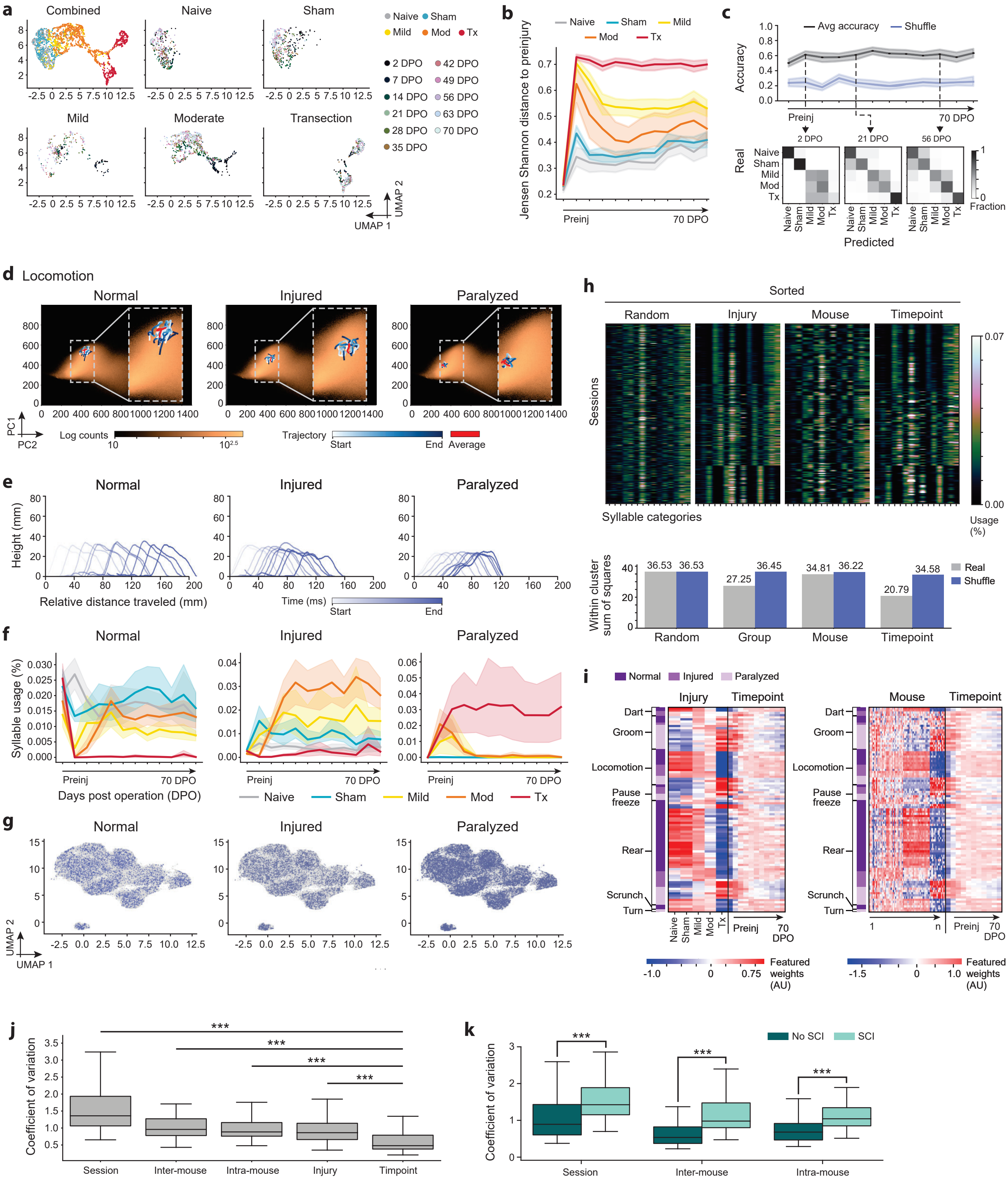

a
Combined
Naive
Sham
Mild
Moderate
Transection
Naive Sham
Mild Mod Tx
2 DPO 42 DPO
7 DPO 49 DPO
14 DPO 56 DPO
21 DPO 63 DPO
28 DPO 70 DPO
35 DPO
UMAP 1
UMAP 2
b
Naive Sham Mild
Mod Tx
Jensen Shannon distance to preinjury
Preinj
70 DPO
c
Avg accuracy Shuffle
Accuracy
Preinj
70 DPO
2 DPO
21 DPO
56 DPO
Real
Naive
Sham
Mild
Mod
Tx
Predicted
Fraction
d Locomotion
Normal
Injured
Paralyzed
PC1
PC2
Log counts
Trajectory
Start
End
Average
e
Normal
Injured
Paralyzed
Height (mm)
Relative distance traveled (mm)
Time (ms)
Start
End
f
Normal
Injured
Paralyzed
Syllable usage (%)
Preinj
70 DPO
Days post operation (DPO)
Naive Sham Mild Mod Tx
g
Normal
Injured
Paralyzed
UMAP 2
UMAP 1
h
Sorted
Random
Injury
Mouse
Timepoint
Sessions
Syllable categories
Usage (%)
Within cluster sum of squares
36.53 36.53
27.25 36.45
34.81 36.22
20.79 34.58
Random
Group
Mouse
Timepoint
Real
Shuffle
i
Normal Injured Paralyzed
Injury
Timepoint
Mouse
Dart
Groom
Locomotion
Pause freeze
Rear
Scrunch
Turn
Naive Sham Mild Mod Tx
Preinj
70 DPO
1
n
Featured weights (AU)
j
Coefficient of variation
***
Session
Inter-mouse
Intra-mouse
Injury
Timpoint
k
No SCI SCI
Coefficient of variation
***
Session
Inter-mouse
Intra-mouse

**Extended Data Fig 2. Injury severity and individual identity jointly structure exhibited behaviors during recovery (female model). a,** UMAPs of syllable usages. Same as Fig. 2a, but for females. **b,** Jensen Shannon distance to preinjury state. Same as Fig. 2b but for females (ANOVA with Dunnett's multiple comparisons test for b and f; n=68; *P* values can be found in Source Data 4). **c,** Linear classifier accuracy in distinguishing groupwise syllable usages across timepoints (top). Example confusion matrices (bottom). Same as Fig. 2c, but for females. **d,** Projection of example locomotion behaviors into PC embedding space. Same as Fig. 2d, but for females. **e,** Spinograms. Same as Fig. 2e but for females. **f,** Longitudinal usage of example locomotion syllable variants. **g,** UMAP embeddings of weighted behavioral sequences, colored by locomotor syllable variants. Same as Fig. 1g, but for females. **h,** Behavioral composition heatmaps sorted randomly or by group, mouse, or timepoint (top). Within cluster sum of squares (bottom). Same as Fig. 1h, but for females. **i,** Weights from generalized linear models with syllable usage as a function of: injury types and timepoints (left) and mouse ID and timepoints (right). Same as Fig. 1i, but for females. **j,** CV of syllable usages across: all sessions (session), different mice (intermouse), sessions within a mouse (intramouse), groups (group), and timepoints (timepoints). Same as Fig. 1j, but for females (for j and k: unpaired t-test; n=68; ***$P < 0.001$; *P* values can be found in Source Data 4). **k,** Same as j, but split by injury (no SCI and SCI) revealing that SCI contributes more to the variation. Same as Fig. 1k, but for females. Mod, moderate; Tx, transection; PC, principal component; UMAP, Uniform Manifold Approximation and Projection; CV, coefficient of variation.

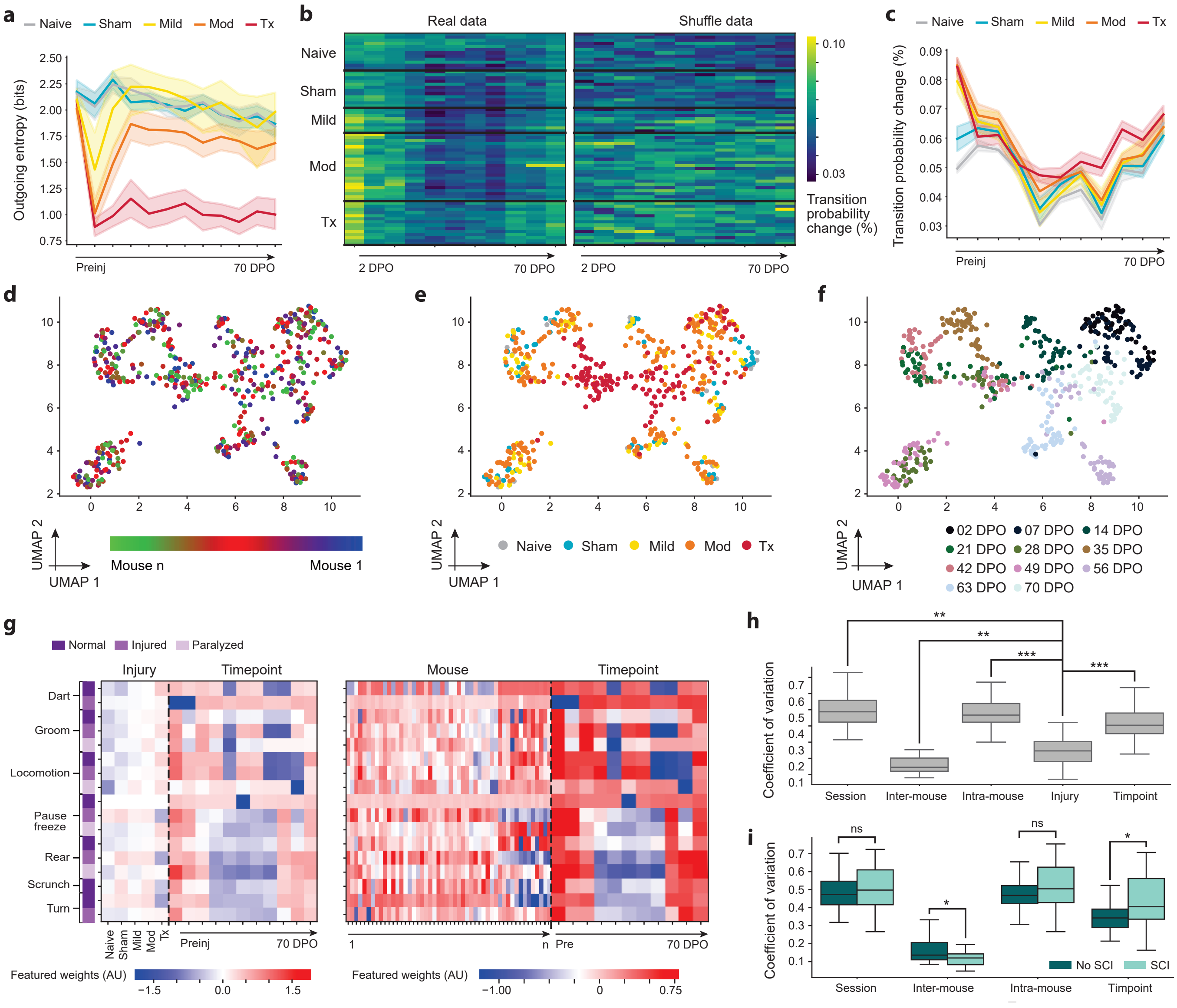


**Extended Data Fig 3. Behavioral sequence reorganization follows a time-dependent structure across recovery (female model). a,** Average syllable outgoing entropy for different injury severities over time (ANOVA with Dunnett's multiple comparisons test for a and c; n=68; *P* values can be found in Source Data 6). Same as Fig. 3b, but for females. **b,** Heatmaps of average absolute changes in outgoing transition probabilities over time using real (left) and shuffle (right) data. Same as Fig. 3c, but for females. **c,** Quantification of average absolute outgoing transition differences in c. Same as Fig. 3d, but for females. **d-f,** UMAP embeddings of transition change vectors across syllable categories, colored by mouse (d), injury severity (e), or timepoint (f). Same as Fig. 3e, but for females. **g,** Weights from generalized linear models relating changes in transition probabilities to injury severity and timepoint (left) or mouse identity and timepoint (right). Same as Fig. 3h but for females. **h,** CV of syllable usages was analyzed across various groupings: session (within all sessions), inter-mouse (across different mice), intra-mouse (within a single mouse), group (across different groups), and timepoints (across different timepoints). Same as Fig. 3i but for females (unpaired t-test; n=68; $*P < 0.05$, $**P < 0.01$, $***P < 0.001$; *P* values can be found in Source Data 6). **i,** Same as i, but split by no SCI vs. SCI. Mod, moderate; Tx, transection; UMAP, Uniform Manifold Approximation and Projection; CV, coefficient of variation.

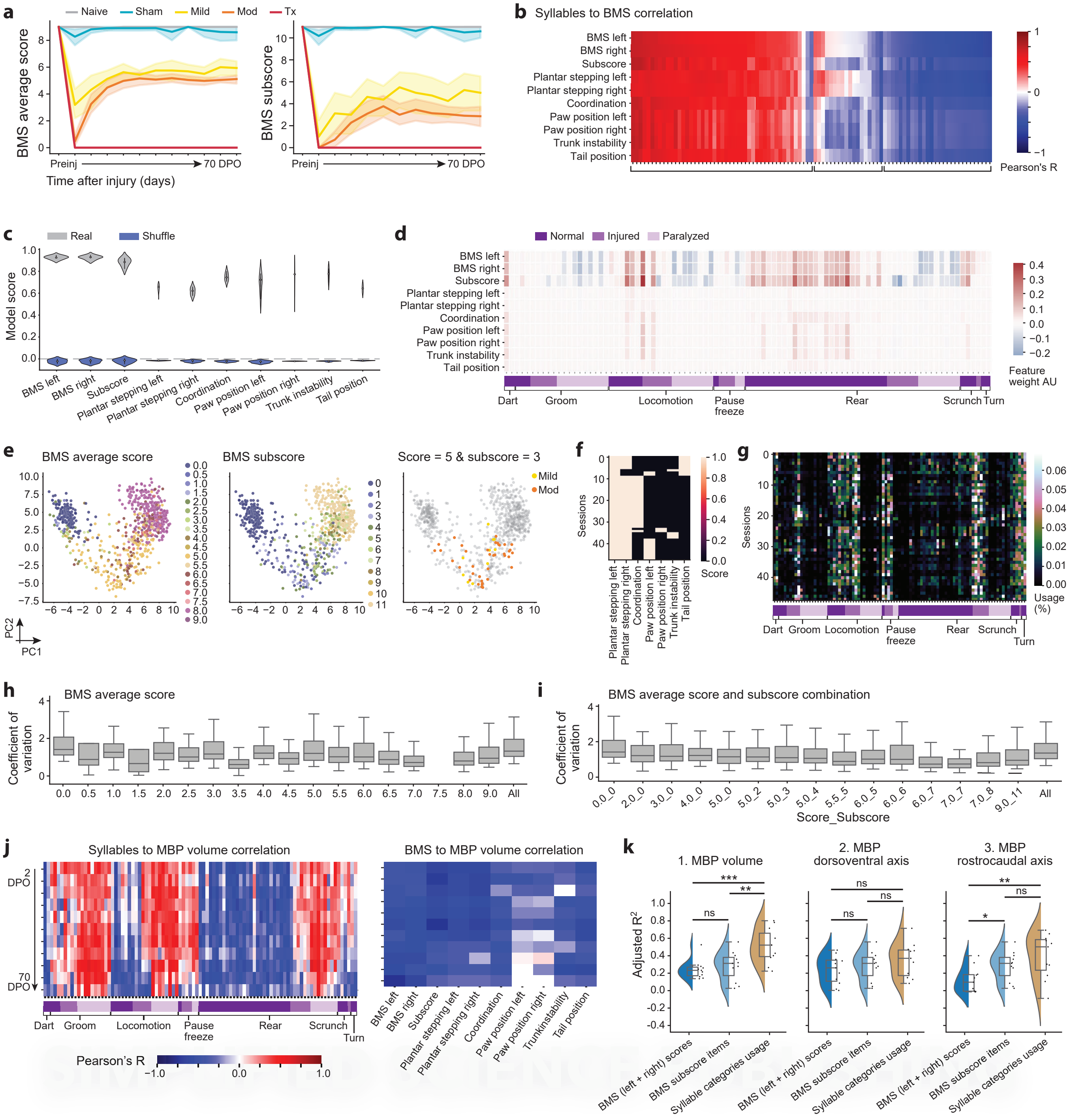
a
Naive
Sham
Mild
Mod
Tx
BMS average score
BMS subscore
Preinj
70 DPO
Time after injury (days)
b
Syllables to BMS correlation
BMS left
BMS right
Subscore
Plantar stepping left
Plantar stepping right
Coordination
Paw position left
Paw position right
Trunk instability
Tail position
Pearson's R
c
Real
Shuffle
Model score
d
Normal
Injured
Paralyzed
Feature weight AU
Dart
Groom
Locomotion
Pause freeze
Rear
Scrunch
Turn
e
BMS average score
BMS subscore
Score = 5 & subscore = 3
Mild
Mod
PC2
PC1
f
Sessions
Score
g
Usage (%)
h
BMS average score
Coefficient of variation
All
i
BMS average score and subscore combination
Score_Subscore
j
Syllables to MBP volume correlation
BMS to MBP volume correlation
2 DPO
70 DPO
Pearson's R
Trunkinstability
k
1. MBP volume
2. MBP dorsoventral axis
3. MBP rostrocaudal axis
Adjusted $R^2$
ns
BMS (left + right) scores
BMS subscore items
Syllable categories usage

**Extended Data Fig 4. Quantitative behavioral descriptions reveal hidden heterogeneity among animals with equivalent locomotor function (female model). a,** BMS average scores (left) and subscores (right) (ANOVA with Dunnett's multiple comparisons test; n = 68; *P* values can be found in Source Data 8). **b,** Pearson's correlation between syllable usages and BMS. Same as Fig. 4b, but for females. **c,** Cross validations of regularized linear regression models predicting BMS criteria from syllable usages. Same as Fig. 4c, but for females. **d,** Feature weights from BMS prediction models in c. Same as Fig. 4d, but for females. **e,** PCA computed from syllable usages colored by BMS scores (left), BMS subscores (middle), and animals with a BMS score-subscore combination of 5-3 (right). Same as Fig. 4e, but for females. **f-g,** Heatmap of BMS criteria for animals with a score-subscore combination of 5-3 (f) and corresponding heatmap of syllable usages for the same animals (g). Same as Fig. 4fg, but for females. **h-i,** CV of syllable usage within individual BMS scores and across all scores (h) and across all BMS score-subscore combinations (i). Same as Fig. 4hi, but for females. **j,** Pearson's correlations for injury-induced demyelination volumes and syllable usages (left) and BMS criteria (right). Same as Fig. 4j, but for females. **k,** Adjusted $R^2$ values from correlation models of lesion metrics with BMS Scores, BMS subscores, and syllable category usages. Same as Fig. 4k, but for females (unpaired t-test; n=68; **P* < 0.05, ***P* < 0.01, ****P* < 0.001; *P* values can be found in Source Data 8). Mod, moderate; Tx, transection; BMS, Basso Mouse Scale; PCA, principal component analysis; MBP, myelin basic protein.

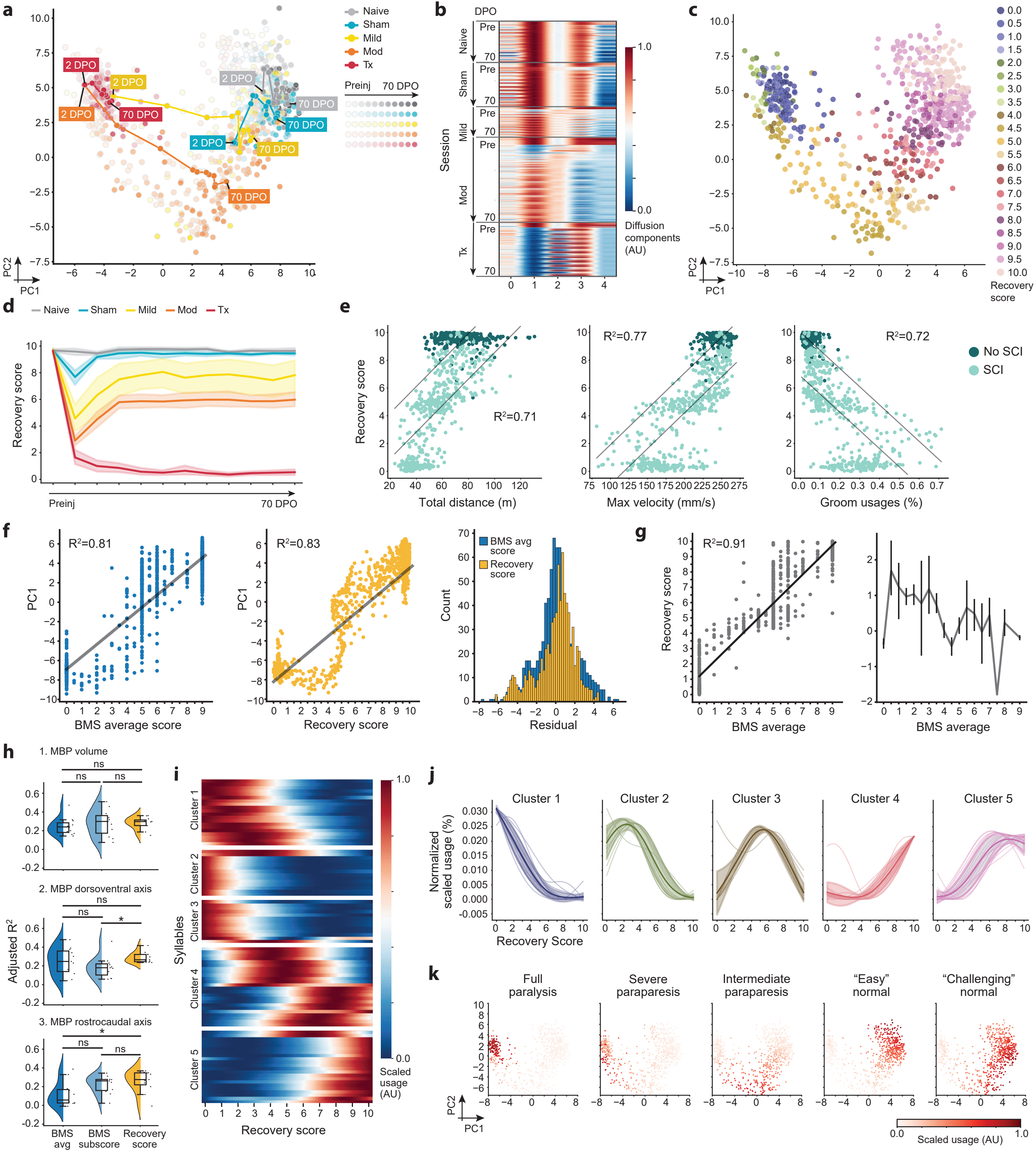

a
Naive
Sham
Mild
Mod
Tx
Preinj
70 DPO
2 DPO
70 DPO
PC2
PC1
b
DPO
Pre
70
Naive
Sham
Mild
Mod
Tx
Session
1.0
0.0
Diffusion components (AU)
c
Recovery score
PC2
PC1
d
Naive
Sham
Mild
Mod
Tx
Recovery score
Preinj
70 DPO
e
Recovery score
R²=0.71
R²=0.77
R²=0.72
Total distance (m)
Max velocity (mm/s)
Groom usages (%)
No SCI
SCI
f
R²=0.81
R²=0.83
PC1
BMS average score
Recovery score
BMS avg score
Recovery score
Count
Residual
g
R²=0.91
Recovery score
BMS average
BMS average
h
1. MBP volume
2. MBP dorsoventral axis
3. MBP rostrocaudal axis
ns
*
Adjusted R²
BMS avg
BMS subscore
Recovery score
i
Cluster 1
Cluster 2
Cluster 3
Cluster 4
Cluster 5
Syllables
Recovery score
1.0
0.0
Scaled usage (AU)
j
Cluster 1
Cluster 2
Cluster 3
Cluster 4
Cluster 5
Normalized scaled usage (%)
Recovery Score
k
Full paralysis
Severe paraparesis
Intermediate paraparesis
“Easy” normal
“Challenging” normal
PC2
PC1
0.0
1.0
Scaled usage (AU)

**Extended Data Fig 5. A repertoire-derived recovery score reveals clusters of coevolving behaviors across the paralysis-to-normality continuum (female model). a,** PCA embedding of syllable usage colored by group, shaded by timepoint, and labeled with each group's averaged longitudinal behavior. Same as Fig. 5a, but for females. **b,** Diffusion components derived from syllable usages to compute the "recovery score." Same as Fig. 5b, but for females. **c,** Projection of recovery scores onto the PCA embedding. Same as Fig. 5c, but for females. **d,** Recovery scores recovery curves (ANOVA with Dunnett's multiple comparisons test; n = 68; *P* values can be found in Source Data 10). Same as Fig. 5d, but for females. **e,** Correlations between recovery scores and total distance traveled, maximum velocity, and groom usages (Ordinary Least Squares linear regression for e-g). Same as Fig. 5e, but for females. **f,** Comparison of linear model fits between behavioral PC1 with BMS scores (left) and recovery scores (middle). Model residuals indicate recovery scores are more uniformaly distributed (right). Same as Fig. 5f, but for females. **g,** Correlations between BMS scores and recovery scores (left) and model residuals (right). Same as Fig. 5g, but for females. **h,** Adjusted $R^2$ values from correlation models relating lesion metrics with BMS scores, BMS subscores, and recovery scores. Same as Fig. 5h, but for females (unpaired t-test; n=68; *$P < 0.05$; *P* values can be found in Source Data 10). **i,** Heatmap of syllable usages across recovery scores. Same as Fig. 5i, but for females. **j,** Normalized syllable usages within each syllable cluster (individual syllables, thin lines; cluster means, bold lines; standard deviation, shading). **k,** PCA embeddings colored by normalized mean cluster-specific syllable usages. Clusters were semantically labeled (Supplemental Table 4). Same as Fig. 5k. but for females. Mod, moderate; Tx, transection; BMS, Basso Mouse Scale; PCA, principal component analysis; MBP, myelin basic protein.
