## Supplemental Figures for "The composition of motor recovery revealed by syllable-level behavioral analysis after spinal cord injury"

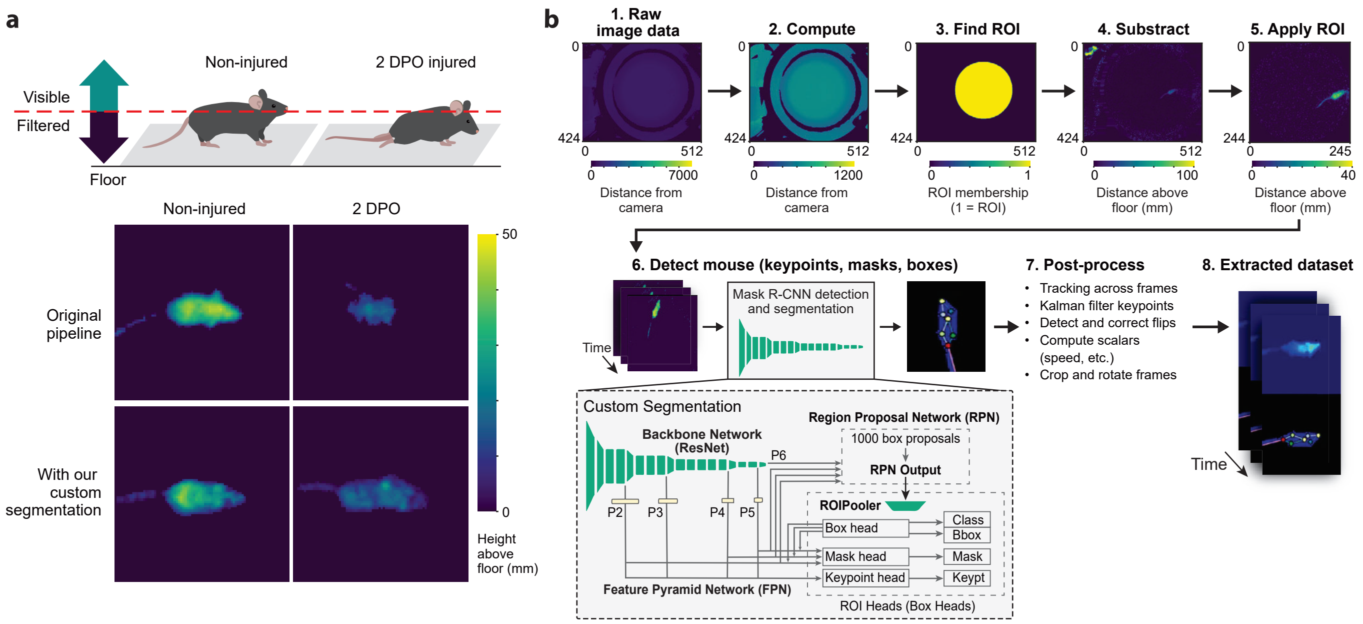


**Supplemental Fig 1. Custom segmentation and syllable analysis for SCI recovery in mice. a,** Illustrative example of the original MoSeq pipeline and our custom segmentation algorithms. The original MoSeq data extraction tools could not support SCI mice as dragging trunks and limbs of injured mice were classified as noise and filtered out due to their significantly lower positioning compared to uninjured mice. We custom adapted the data processing step by leveraging the Detectron2 implementation of multi-task RCNN for appropriate full body segmentation of SCI mice. **b,** Schematic of the extraction pipeline, including the addition of our custom segmentation algorithms. DPO, days post operation; MoSeq, motion sequencing; ROI, region of interest; RCNN, region convolutional neural network. Custom segmentation code can be found on our GitHub: https://github.com/tischfieldlab.

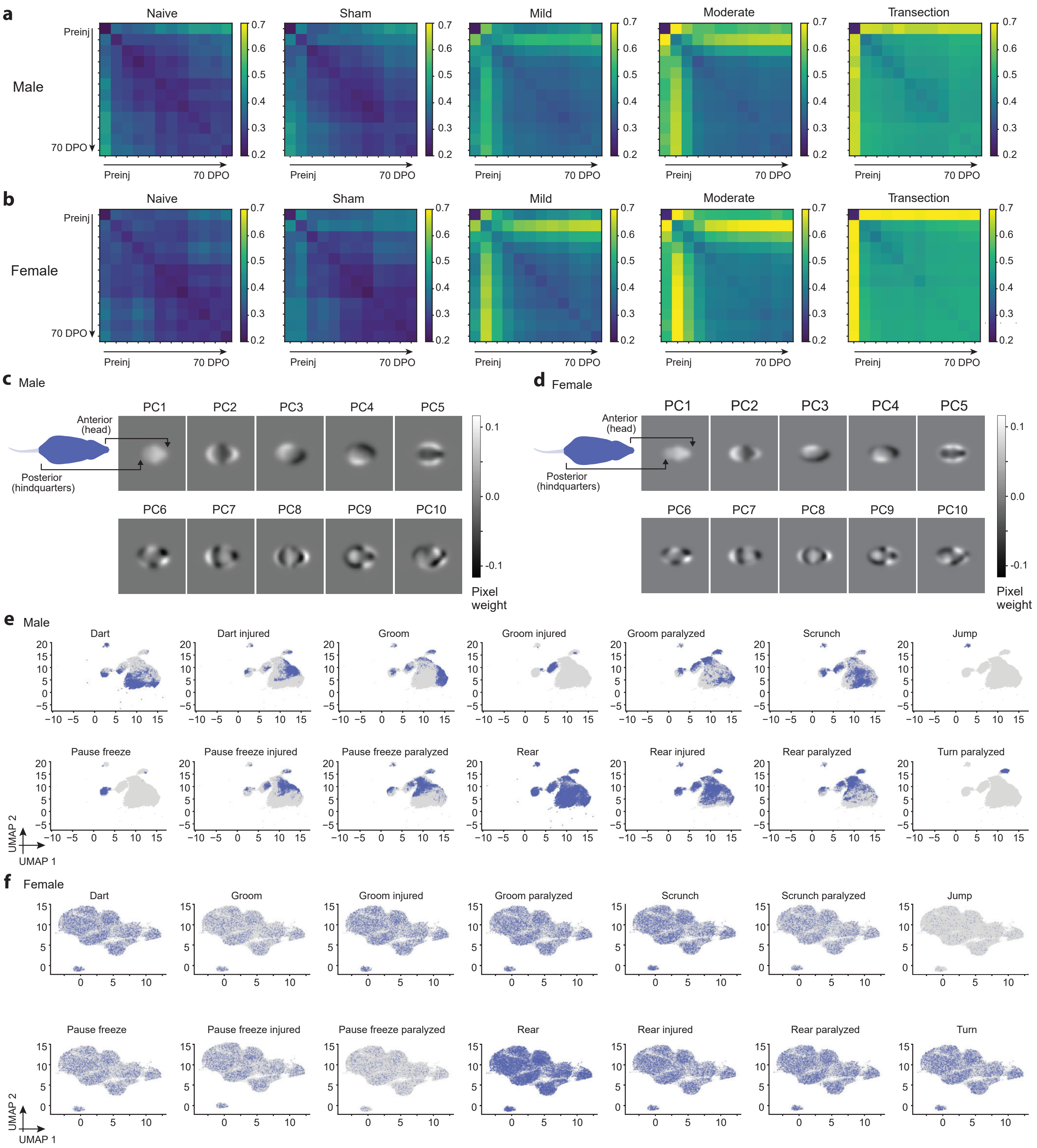


**Supplemental Fig 2. Pose dynamics and sequence structure underlying behavioral composition identification. a,** Heatmap plots for each group of the Jensen Shannon distance to preinjury timepoint over time for the male model, denoting how similar timepoints post operation are to the preinjury timepoint. **b,** Same as a, but for the female model. **c,** The top 10 PCs, used by the AR-HMM model to capture pose dynamics and segment syllables, illustrate distinct pixel weight distributions. For example, PC1 primarily reflects variations in head height. **d,** Same as c, but for the female model. **e,** UMAP plots of weighted syllable sequences, colored by all syllable categories (except for locomotion syllables) across different mobilities for the male model, demonstrate that structural patterns vary distinctly between syllables associated with different mobility states. **f,** Same as c, but for the female model. Mod, moderate; Tx, transection; PC, principal component; AR-HMM, autoregressive hidden Markov model; UMAP, Uniform Manifold Approximation and Projection.

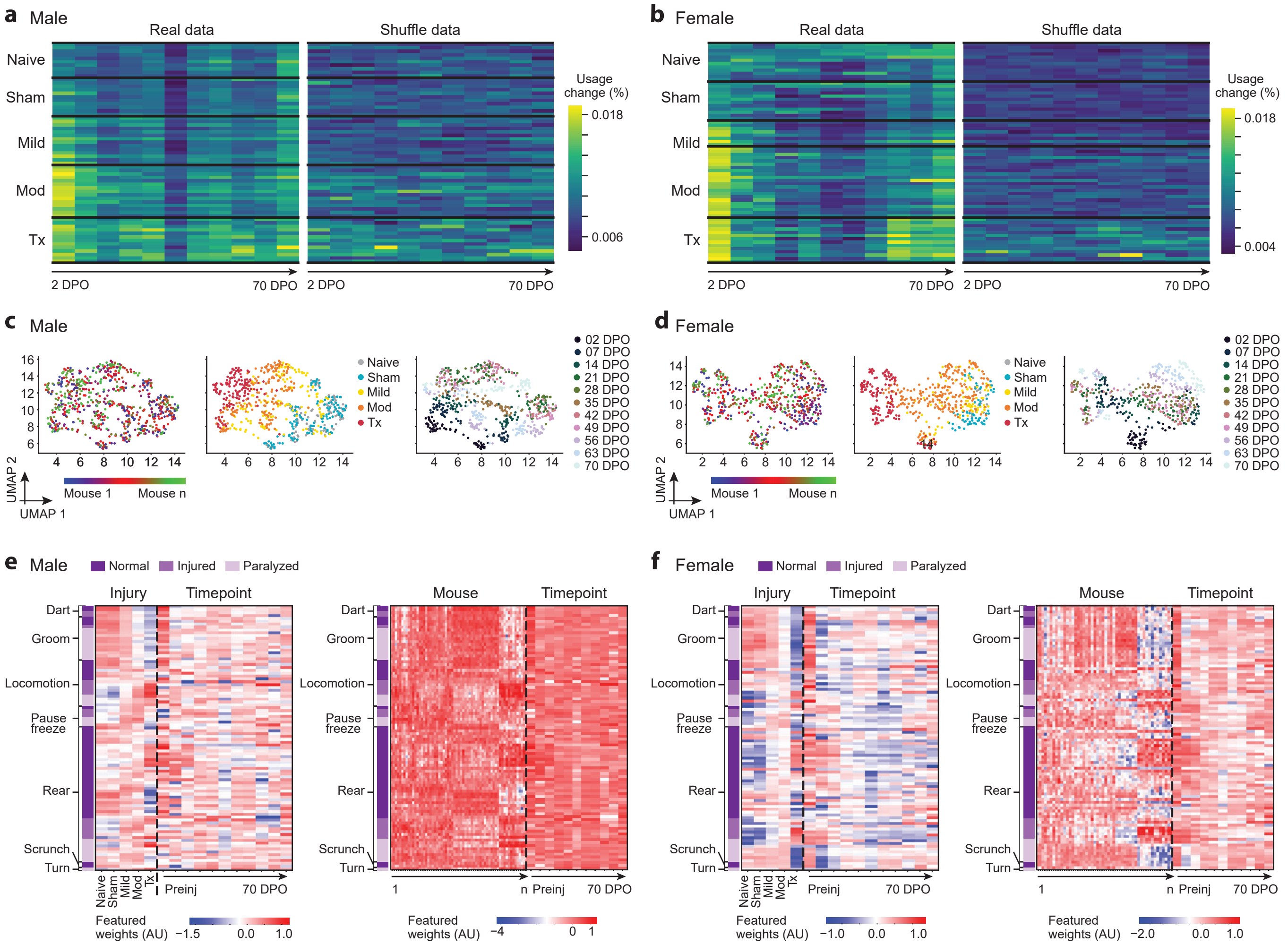


**Supplemental Fig 3. Temporal dynamics and variability in syllable usage differences across groups and timepoints. a,** Average absolute syllable usage differences of 17 syllable categories heatmap over time using real (left) and shuffle (right) data. **b,** Same as c, but for the female model. **c,** UMAP of average absolute syllable usage differences of 17 syllable categories for the male model, colored by mouse (left), group (middle), and timepoint (right) (see methods). **d,** Same as e, but for the female model. **e,** Weights from generalized linear models for the male model with average absolute syllable usage differences as a function of two factor types: injury types and timepoints (left) and mouse and timepoints (right). **f,** Same as g, but for the female model. Mod, moderate; Tx, transection; UMAP, Uniform Manifold Approximation and Projection.

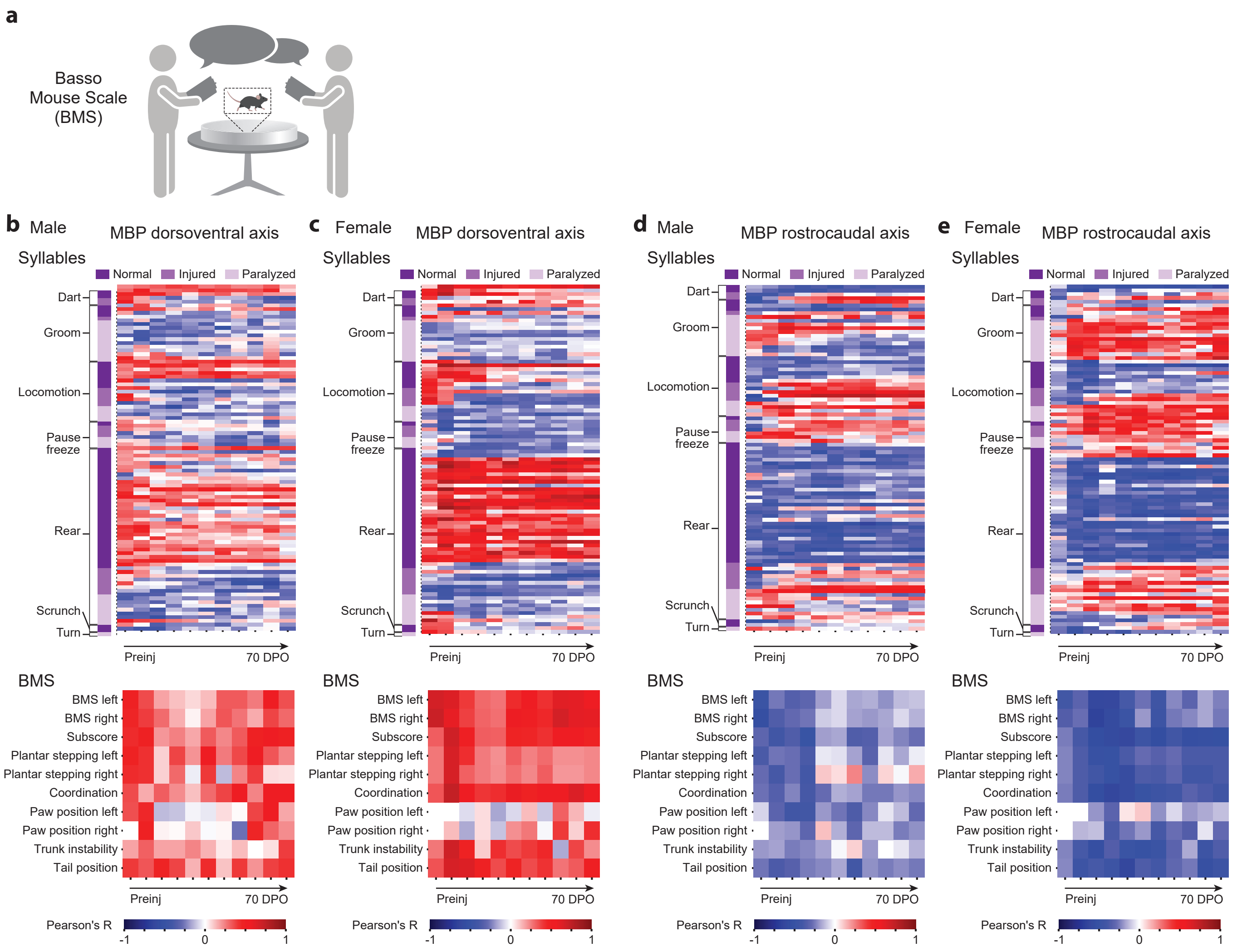


**Supplemental Fig 4. Correlations with injury-induced demyelination across injury axes. a,** Illustrative schematic of manual BMS scoring conducted by two trained evaluators. **b,** Pearson's correlations for injury-induced demyelination along the dorsoventral axis and syllable usages (top) and BMS criteria (bottom) for the male model. **c,** Same as a, but for the female model. **d,** Pearson's correlations for injury-induced demyelination along the rostrocaudal axis and BMS criteria (top) and syllable usages (bottom) for the male model. **e,** Same as c, but for the female model. Mod, moderate; Tx, transection; BMS, Basso Mouse Scale; MBP, myelin basic protein.

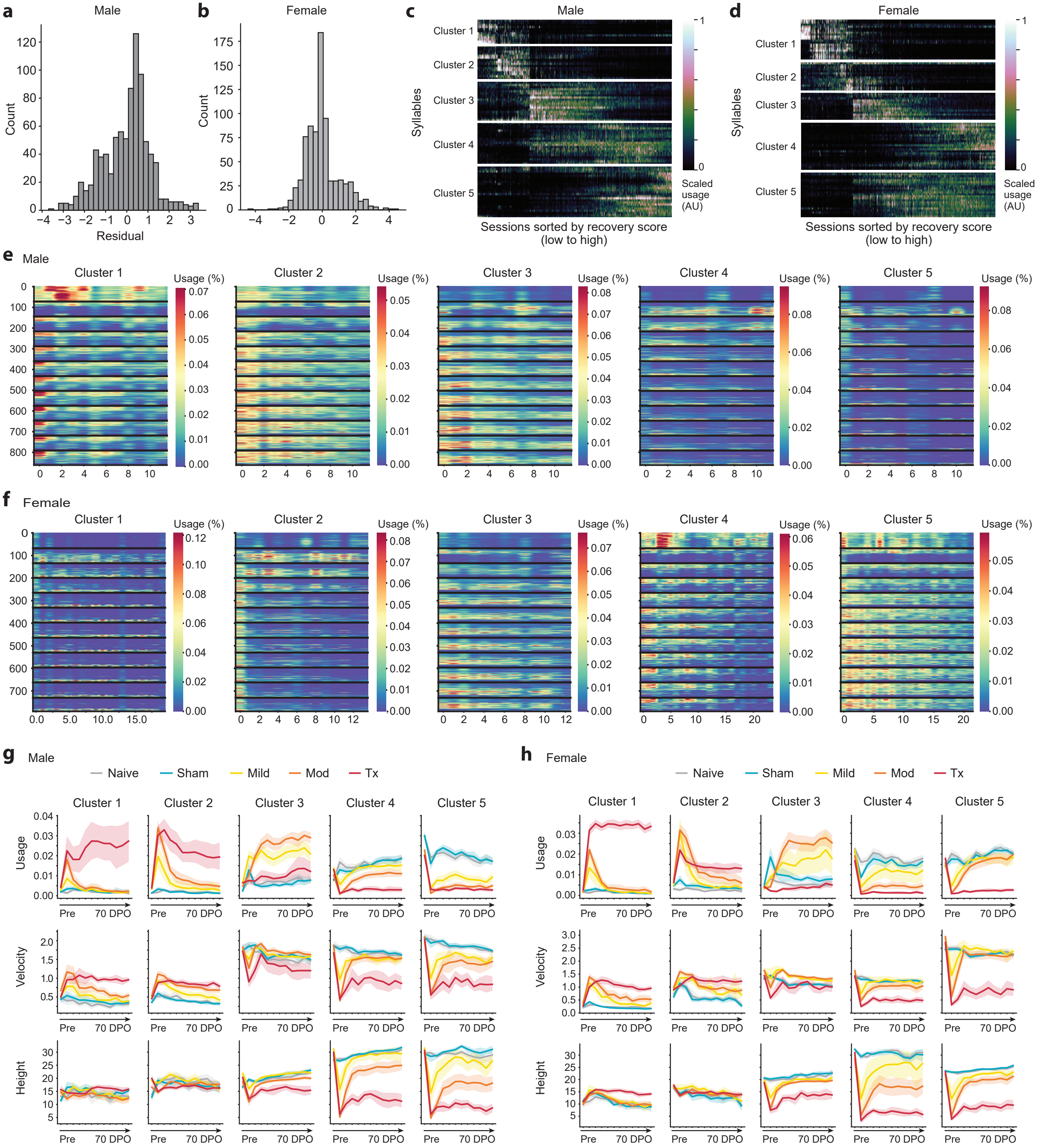


**Supplemental Fig 5. Trends of syllable usages across Recovery Scores and syllable clusters.** **a,** Residual values histogram for the correlation of BMS average scores and recovery scores for the male model. **b,** Same as a, but for the female model. **c,** Heatmap of raw syllable usage trends calculated using the recovery score model (see methods) for the male model. **d,** Same as c, but for the female model. **e,** Heatmap plots of the normalized usage of syllables within each cluster for the male model. **f,** Same as e, but for the female model. **g,** Line plots of usages (top), velocity (middle), and height (bottom) by group over time for each cluster for the male model (n = 71; ANOVA with Dunnett's multiple comparison test here and below; *P* values can be found in Source Data 11). h, Same as g, but for the female model (n = 68, Source Data 11).
